# Defining the transcriptional responses of *Aspergillus nidulans* to cation/alkaline pH stress and the role of the transcription factor SltA

**DOI:** 10.1101/2020.04.16.044396

**Authors:** Irene Picazo, Oier Etxebeste, Elena Requena, Aitor Garzia, Eduardo A. Espeso

**Affiliations:** Department of Cellular and Molecular Biology, Centro de Investigaciones Biológicas Margarita Salas, CSIC. Ramiro de Maeztu, 9, 28040 Madrid, Spain; Laboratory of Biology, Department of Applied Chemistry, Faculty of Chemistry, University of The Basque Country, 20018 San Sebastian, Spain; Laboratory of RNA Molecular Biology, Rockefeller University, New York, USA; Department of Plant Protection, Instituto Nacional de Investigación y Tecnología Agraria y Alimentaria, Ctra. de La Coruña Km. 7, 28040 Madrid, Spain

**Keywords:** Fungi, *Aspergillus nidulans*, cation stress, alkaline pH stress, signal transduction, transcriptional control, SltA, PacC/Rim101

## Abstract

Fungi have developed the ability to overcome extreme growth conditions and thrive in hostile environments. The model fungus *Aspergillus nidulans* tolerates, for example, ambient alkalinity up to pH 10 or molar concentrations of multiple cations. The ability to grow under alkaline pH or saline stress depends on the effective function of, at least, three regulatory pathways mediated by high hierarchy zinc-finger transcription factors: PacC, which mediates the ambient pH regulatory pathway, the calcineurin-dependent CrzA and the cation-homeostasis responsive factor SltA. Using RNA sequencing, we determined the effect of external pH alkalinisation or sodium stress on gene expression. Data show that each condition triggers transcriptional responses with a low degree of overlap. By sequencing the transcriptomes of the null mutant, the role of SltA in the abovementioned homeostasis mechanisms was also studied. Results show that the transcriptional role of SltA is wider than initially expected and implies, for example, the positive control of the PacC-dependent ambient pH regulatory pathway. Overall, our data strongly suggest that the stress-response pathways in fungi include some common but mostly exclusive constituents, and that there is a hierarchy of authority among the main regulators of stress response, with SltA controlling *pacC* expression at least in *A. nidulans*.

## Background

The survival of organisms is subject to their ability to adapt to and overcome diverse challenges or stresses in the environment. The most general responses are produced by coordinating the regulation of gene expression and those mechanisms controlling the proper localization of proteins in the cell. Gene expression can be enhanced or repressed by the activity of transcription factors, which recognise specific consensus sequences at gene promoters through their DNA-binding domains. Concurrently, the activity of a transcription factor can be modulated by other regulatory elements or by post-translational modifications which may occur via elements belonging to a signaling cascade activated by a given ambient stimulus.

The model organism *Aspergillus nidulans* is capable of surviving to a variety of ambient stresses such as wide range of pH values or high extracellular concentrations of ions [1]. Tolerance to ambient alkalinity has been largely studied in this fungus [2–4]. Previous work has led to the identification and functional characterization of three zinc-finger transcription factors that coordinate a response to ambient alkaline pH: PacC, regulator of the ambient pH regulatory pathway [2]; the calcineurin-dependent transcription factor CrzA and the cation-homeostasis transcription factor SltA [5]. The absence of any of these transcription factors inhibits colonial growth of *A.nidulans* at alkaline pH [5].

PacC is a 674 amino acid transcription factor having an N-terminally located DNA-binding domain consisting of three classical Cys2His2 zinc-fingers and C-terminally located key regulatory regions required for the signalling function of this transcription factor [2, 6–8]. PacC can be found in three forms in the cell depending on extracellular pH conditions [9]. At extreme acid pH (4 or lower), the full-length form of 72kDa is mainly detected, which is in a close conformation to prevent its proteolysis [8]. At extracellular alkaline pH, PacC is activated through two proteolytic steps. The first step is pH-dependent and signalled by the Pal pathway [9]. PalH is the pH sensor protein at the plasma membrane, and is stabilized by PalI, another membrane-embedded protein [10]. At alkaline pH, PalF, an arrestin-like protein, becomes phosphorylated and ubiquitylated, and then binds to the C-terminal region of PalH [11]. The PalH-PalF interaction allows the recruitment by PalA to PacC of the rest of Pal-pathway elements: Vps32, PalC and PalB [11–13]. Then, the putative calpain-protease PalB cleaves the PacC^72kDa^ form at a specific site rendering the intermediate form of PacC^53kDa^ [14]. The second proteolytic step is pH-independent, carried out by the proteasome, resulting in the active form of 27kDa [15]. PacC plays a dual function at alkaline pH by repressing those genes expressed at acid pH (acid-expressed genes) and positively regulating those genes whose function is required at alkaline pH (alkaline genes) [16, 17]. An example of this dual function is the opposed regulation of acid and alkaline phosphatases [18] or xylanases [19]. This pH-dependent regulation of gene expression is possible due to the direct binding of PacC to 5’-GCCARG-3’ sequences at target promoters [2, 16, 17].

In contrast to PacC and CrzA, which can be found in most Ascomycetes, SltA is a transcription factor exclusive of Pezizomycotina subphylum [5, 20]. Only two elements have been described in the Slt pathway so far: SltA and the signaling protein SltB, also specific of Pezizomycotina [21]. SltA is a 698 amino acid protein with three classical Cys_2_His_2_ zinc-fingers located between residues 416 and 500, and binding to the 5’-AGGCA-3^1^ target sequence [5, 22]. Initially, *sltA* gene was identified by studying the loss-of-function mutation *sltA1*, which truncates the protein at amino acid 502 and confers *A. nidulans* sensitivity to NaCl, UV radiation, MNNG, 4NQO and arginine [23]. Further studies showed that absence of SltA function causes sensitivity to elevated extracellular concentrations of other cations such as potassium, lithium, caesium or magnesium, but not to calcium, and extreme sensitivity or complete inhibition of colonial growth under ambient alkaline pH [5]. Similarly to PacC, SltA can be simultaneously found in the cell in three forms: a full-length form of 78 kDa and two versions of the proteolysed form of 32kDa, one of which is phosphorylated [20]. This single proteolytic step is driven by the 1272 amino acid protein SltB, a serine protease with a large prodomain similar to a pseudo-kinase domain [20, 21]. SltB is essential for SltA proteolysis but also SltA transcriptional activity is necessary for *sltB* expression [20].

Despite the abundant genetic and molecular knowledge on these regulatory systems, sparse data is available on the transcriptional response of *A. nidulans* to ambient pH alkalinisation or the stress caused by an elevated concentration of sodium. When cultured on solid medium, an inhibition of asexual sporulation at pH 8 and less dense growth in high sodium are observed in the wild-type strain, indicating differential effects of these stresses in colonial morphology (Figure 1A). As reported previously [5, 20], the absence of SltA function causes minor defects in colony growth on standard minimal medium but strong sensitivity to both alkaline pH and high sodium stresses.

**Figure 1:**
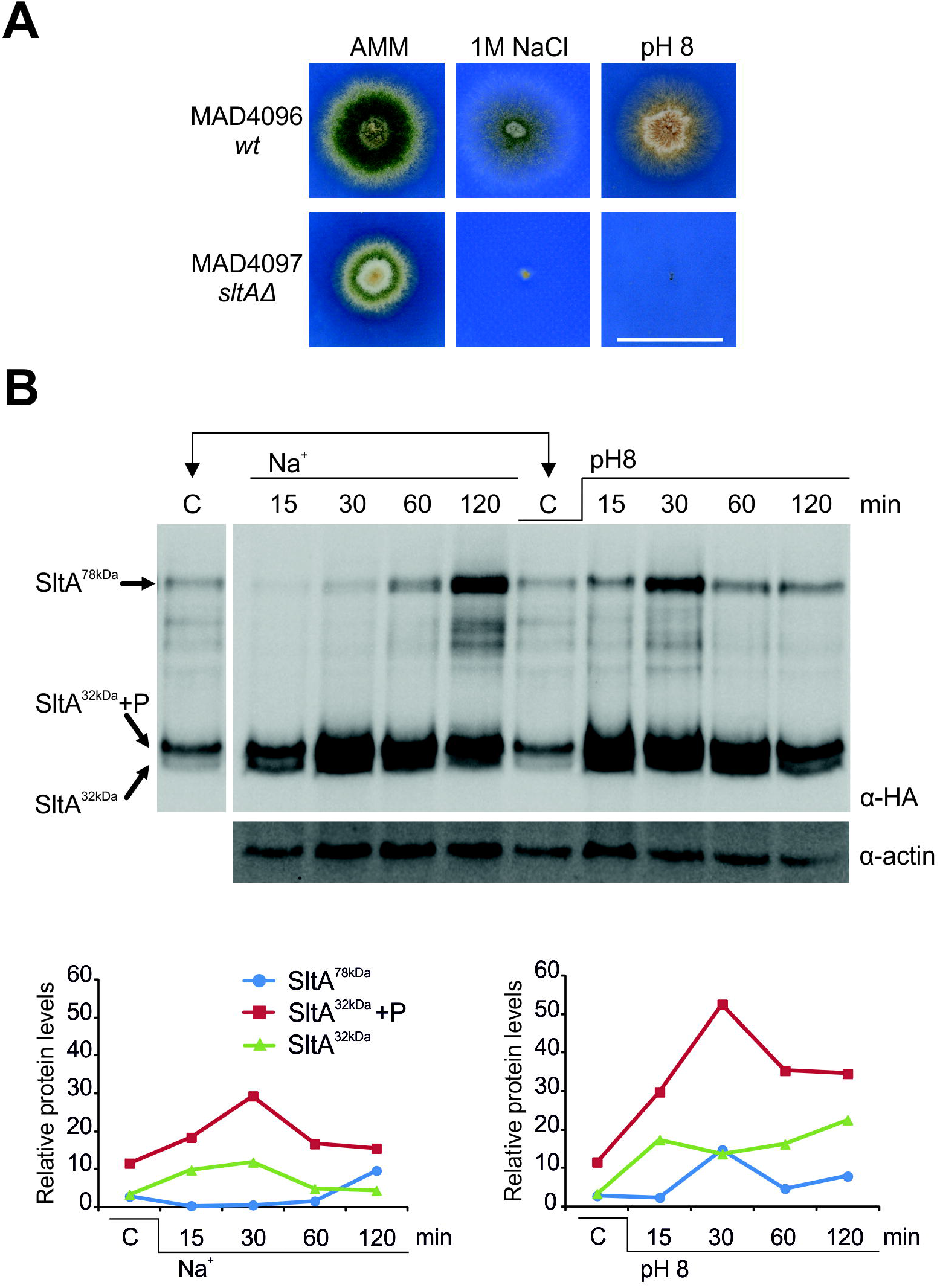
Phenotype of the *sltA*Δ strain and levels of the different SltA forms under cation or alkaline pH stresses. A) Colonial growth of wild-type (WT) and *sltA*Δ strains on AMM, AMM supplemented with 1.0 M NaCl or an alkalinised medium (addition of 100 mM Na_2_HPO_4_; pH 8), after 48 hours of culture at 37 °C. Scale bar = 2 cm. B) Immunodetection of SltA::HA_3_ in protein extracts obtained from mycelial samples grown in AMM (C: control), or at 15, 30, 60 and 120 minutes after the addition of 1.0 M NaCl or 100 mM Na_2_HPO_4_ (pH 8). α-actin was used as control and to quantify the levels of the different SltA forms detected. Bottom: Charts showing the relative levels of SltA forms. Values were obtained by calculating the ratio between SltA and α-actin band intensitites.

To determine the set of genes up- or downregulated under the abovementioned stress conditions, the role of SltA in the corresponding homeostasis mechanisms and the existence of a transcriptional dependence of the other two high-hierarchy regulators (or other unknown transcription factors) on SltA, we carried out RNA sequencing experiments of wild-type and *sltAΔ* samples followed by multiple comparisons of the transcriptomes. Our results show that there is a low degree of overlap between the transcriptional responses of *A. nidulans* to medium alkalinisation or sodium stress, suggesting that each stress-response pathway includes mostly exclusive constituents. In both cases, the expression of specific subsets of genes remains independent of SltA activity. Other subsets positively depend on SltA, including the PacC-dependent ambient pH regulatory pathway, suggesting that there is a hierarchy of authority among the main regulators of stress response in fungi.

## Methods

### Strains, oligonucleotides and culture conditions

Oligonucleotides used in this study are shown in Supplementary Table S1 while the genotype of the strains used is shown in Supplementary Table S2. Strain MAD6669 was obtained by transformation of strain MAD1427 using a fusion PCR fragment as described before [20], allowing the tagging of SltA with three copies of hemagglutinin epitope (HA_3_). The construct contained the *riboB* gene from *Aspergillus fumigatus* (*riboB^Af^*), which was used as the selection marker in the transformation.

Sensitivity of the null *sltA* mutant to sodium stress or medium alkalinity was compared to that of the reference wild-type strain, by culturing both strains on adequately supplemented solid Aspergillus minimal medium or AMM [24], medium supplemented with 1.0 M NaCl (sodium stress) or a medium alkalinised to pH 8 by adding 100 mM Na_2_HPO_4_ [5, 21].

Samples for RNA extraction were cultured as follows (see also diagram in Figure 2A). Conidia (10^6^ conidia/mL) of each of the strains were inoculated into 100 mL Erlenmeyer flasks (twelve per strain, four sets of biological samples) containing 25 mL of adequately supplemented liquid standard AMM. After 18 hours of culture at 37 ºC and 250 rpm, for each set, NaCl (1.0 M final concentration) or Na_2_HPO_4_ (100 mM final concentration) were added, and one of the flasks of each set was kept as control. Cultures were further incubated for 60 minutes. Then, mycelia were collected and two sets were combined to minimize variations. pH of filtered media was monitored and both control and NaCl samples showed an approximate final pH value of 4, and pH 8 the sample containing Na_2_HPO_4_. Mycelia of two combined samples were processed as duplicates.

**Figure 2:**
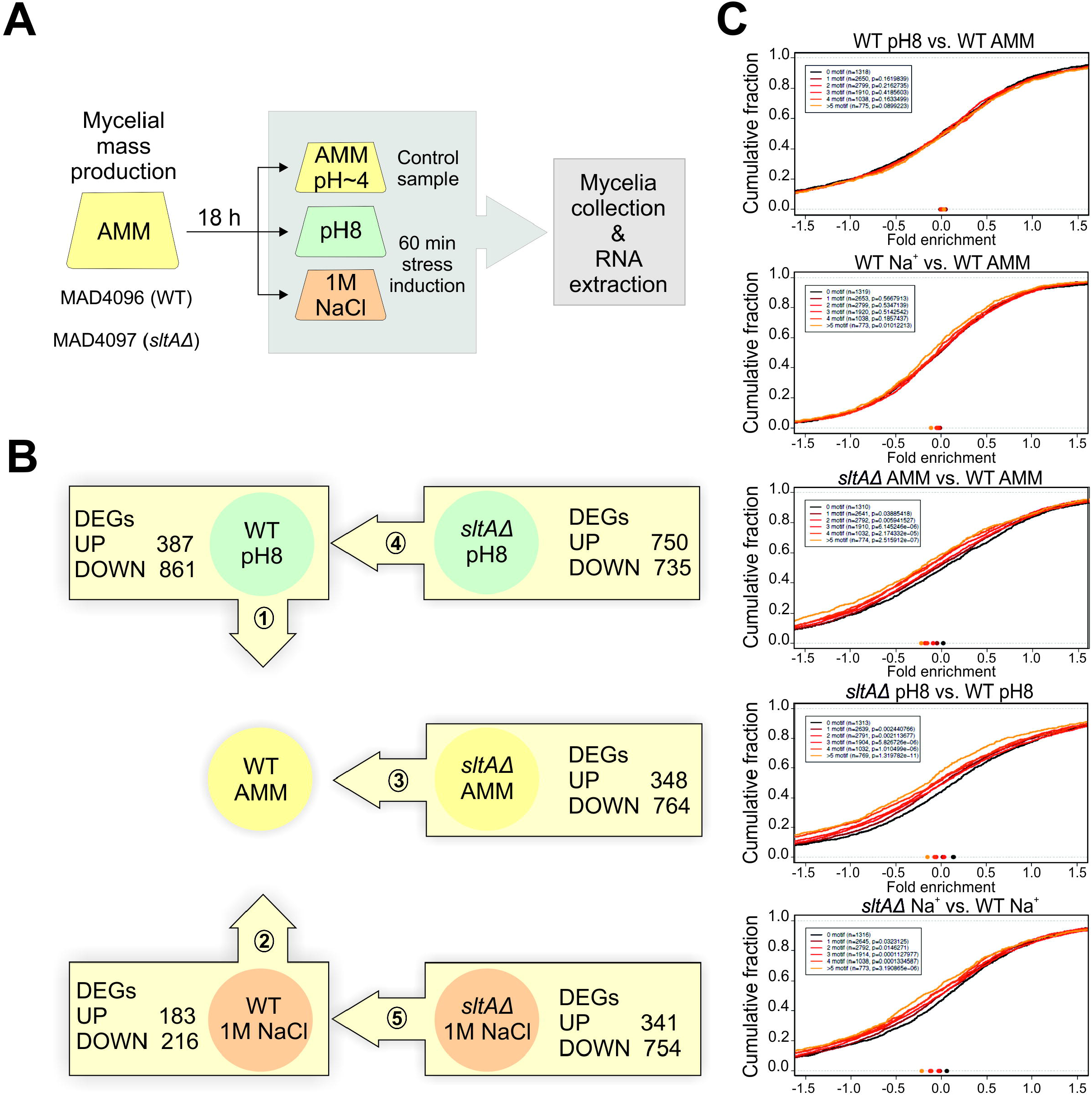
Experimental design and number of significantly de-regulated genes in each RNA sequencing comparison. A) Experimental design for the RNA sequencing experiment carried out in this work. Spores of wild-type and *sltA*Δ strains were collected and incubated (10^6^ conidia/mL) in AMM for 18 hours at 37 °C. Then, Na_2_HPO_4_ (100 mM) or NaCl (1.0 M), final concentrations in both cases, were added, followed by an incubation of 60 minutes. Cultures grown in AMM were used as controls. Samples (duplicates; see Materials and Methods details) were then filtered and processed for RNA extraction and sequencing. B) Diagram representing the five transcriptional comparisons carried out in this work. Arrows indicate the direction of the comparison and boxed are the number of significantly up- and downregulated genes in each transcriptional comparison. C) Cumulative-distribution fraction plots correlating, in each of the five transcriptomic comparisons, log2FC values with the number of SltA target sites in gene promoters. Groups of promoters (1,000 bp-s upstream of the starting ATG) with 0, 1, 2, 3, 4 and >5 SltA target motifs were differentiated and plotted. The points in the lower part of each graph indicate when 50% of the genes are reached. The value of p for each plot is included.

Samples for protein extraction were cultured as above in liquid AMM for 18 hours. After the addition of 1.0 M NaCl or 100 mM Na_2_HPO_4_, mycelial samples were collected at 15, 30, 60 and 120 minutes and processed according to the alkaline lysis procedure (see below; [25]).

### RNA extraction

Mycelia cultivated under the conditions described in the previous section were harvested by filtration using Miracloth (Calbiochem, Merck-Millipore, Darmstadt, Germany). Mycelial samples of approximately 300 mg were frozen and ground in liquid nitrogen before the addition of 1 mL per sample of TRIreagent (Fluka, Sigma-Aldrich Quimica SL, Madrid, Spain) and further processing according to the manufacturer’s protocol. After TRIreagent addition, samples were incubated for 5 minutes at RT. Then, chloroform (0.2 mL per sample) was added, mixed vigorously, incubated for 15 minutes at RT and centrifuged at 12,000 g and 4 °C during 15 minutes. After collecting the aqueous (upper) phase, 2-propanol (0.5 mL per sample) was added to precipitate RNA, followed by a new centrifugation at 12,000 g and 4 °C during 10 minutes. Ethanol (75%; 1 mL) was added to wash RNA, followed by a centrifugation at 7500 g and 4 ºC during 5 minutes., air drying and addition of 200 μL of RNA-free milliQ water. A duplicate RNA extraction was done for each mycelial sample. Total RNA samples were frozen at −80 °C until sequencing or use in quantitative PCR (qPCR).

### Library construction and RNA sequencing

Library construction and RNA sequencing were carried out in Stabvida (Caparica, Portugal). Quality control of the total RNA samples was based on an RNA integrity number (RIN) equal to or higher than 6.5. Samples (input of ≥ 1 μg total RNA, free of contaminating DNA) were directed for library preparation using Kapa Stranded mRNA Library Preparation Kit (Roche Diagnostics Corporation, Indiana, USA) and sequencing in an Illumina HiSeq 4000 platform, using 150bp paired-end sequencing reads. RNA sequencing data was deposited in the NCBI Sequence Read Archive (SRA) under BioProject ID PRJNA625291.

### Bioinformatic analyses

Adapters were trimmed using cutadapt, version 2.5 (https://cutadapt.readthedocs.io/en/stable/; [26]). Reads were aligned to an Aspergillus genome reference file downloaded from the FungiDB database (FungiDB_36_AnidulansFGSCA4; [27]) using STAR aligner (version 2.5.2a; [28]) and allowing no more than two mismatches. Counts for genes were generated using the featureCounts (version 1.4.6; [29]) in the Rsubread package. The TPM calculation was performed using Rsem (version 1.2.31; [30]) while sample distance or PCA plots, as well as differential expression analyses were carried out using DESeq package [31]. Cumulative-distribution fraction plots were generated using R (version 3.3.3; [32]).

Domain analyses of protein sequences were done using Interpro [33]. Lists of DEGs were initially analyzed using the FungiDB database (https://fungidb.org/fungidb/), including the distribution of DEGs by chromosomes, or the presence of signal peptides and/or transmembrane domains in the corresponding polypeptides. Fungifun (https://elbe.hki-jena.de/fungifun/) was used for GO enrichment analyses [34] while Interpro domain-enrichment analyses were carried out in the ShinnyGO website (http://bioinformatics.sdstate.edu/go/; [35]). Heatmaps showing the expression changes of DEGs were generated with Heatmapper (http://www.heatmapper.ca/; [36]), using average linkage clustering and Pearson distance measurement methods. Expression clusters were inferred and the list of genes in each one was obtained also from Heatmapper. GraphPad was used to plot the expression pattern of genes in each cluster as well as ShinnyGO and Fungifun results, and to obtain the correlation coefficients between RNA sequencing duplicates. Venn diagrams were generated and downloaded with InteractiveVenn (http://www.interactivenn.net/; [37]).

### Quantitative PCR

The SuperScript First-strand Synthesis System for RT-PCR (Invitrogen) was used to synthesize cDNA, following manufacturer’s instructions and after having eliminated traces of genomic DNA (DNAg) using Deoxyribonuclease I, amplification grade (Invitrogen). Two micrograms of total RNA were treated with 1 unit of DNAseI (Invitrogen) for 15 minutes at RT (reaction volume of 10 μL). DNAg digestion was stopped by addition of 1 μL of EDTA and incubation for 10 minutes at 65 °C. Total RNA concentration in samples was determined using a NanoDrop ND-1000. First strand cDNA reaction consisted of 1 μg of DNAseI-treated RNA, 1 μL of 10 mM dNTPs mixture, 1 μL of oligonucleotide oligo(dT)12–18 and milliQ water up to a final volume of 10 μL. Samples were incubated at 65 °C and during 5 minutes and then immediately transferred into ice for 1 minute. After ice incubation, a mixture consisting of 2 μL of RT buffer 10X, 4 μL of MgCl_2_ 25 mM, 2 μL of DTT 0.1 M and 1 μL of the enzyme RNAseOUT™ (40 U/L) was added. After centrifugation at maximum speed for 20 s, samples were incubated at 42 °C for 2 minutes. Then, 1 μL of the retrotranscriptase SuperScript™ II RT was added to each reaction and incubated at 42 °C for 50 minutes, followed by a second incubation at 70 °C for 15 minutes, allowing it to cool on ice for 2–3 minutes at the end of the process. Finally, 1 μL of the enzyme RNAseH was added per sample and the samples were incubated at 37 °C during 20 minutes. The amount of cDNA obtained was measured in a NanoDrop ND-1000. Samples were stored at −20 °C until use.

Once the cDNA samples of the strains of interests were generated, dilutions were made at the final concentrations of 100 ng/μL and 10 ng/μL in water PCR Probe. Each reaction mixture contained 1 μL of the corresponding dilution, 0.3 μL (300 nmoles) of each of the relevant oligonucleotides (Supplementary Table S1), 8.4 μL of PCR Probe water and 10 μL of the 2x commercial reaction mixture (SYBR^®^ Green Master Mix, Bio-Rad. A LightCycler96^®^ (Roche) device was used and PCR conditions applied were: pre-incubation at 95 °C/5 minutes and 40 cycles of 95 °C/10 seconds + 65 °C/30 seconds. To obtain the melting curve, a cycle was used in which the temperature was lowered from 95 °C to 65 °C at a speed of 4.4 °C every 10 seconds.

### Protein extraction and immunodetection

Protein extracts for immunodetection were obtained through the alkaline lysis protocol [25]. Briefly, approximately 6 mg of lyophilized mycelium were resuspended in 1 mL lysis buffer (0.2M NaOH, 0.2 % β-mercaptoethanol). After trichloroacetid acid (TCA) precipitation, 100 μL Tris-Base (1 M) and 200 μL of loading buffer (62.5 mM Tris-HCl pH=6.8, 2 % SDS (p/v), 5 % β-mercaptoethanol (v/v), 6 M urea and 0.05 % bromophenol blue (p/v)) were added. Samples were then loaded and proteins separated on SDS-polyacrylamide gels (%10), and electro-transferred to nitrocellulose filters (Trans Blot Turbo transfer packs) using the Trans Blot Turbo system (BioRad) following the manufacturer’s instructions. HA_3_-tagged proteins were detected using a monoclonal rat α-HA3 (1:1000; clone 3F10; Sigma-Aldrich). Actin, used as loading control, was detected using monoclonal mouse α-actin (1:5000; clone C4; MP Biomedicals). Peroxidase conjugated goat anti-rat IgG+IgM (1:4000; Southern Biotech) or anti-mouse (1:4000; Jackson ImmunoResearch Laboratories) were used as secondary antibodies. Western blots were developed using Amersham Biosciences ECL kit and chemiluminescence was detected using a Chemidoc™ image system driven by Image Lab™ Touch software (version 2.2.0.08; BioRad). Images were processed to a minimum using Image Lab™ software (version 6.0; BioRad).

## Results

### Modification of the proteolytic pattern of SltA in response to sodium or alkaline pH stress

To establish the timing for sample collection, immunodetection experiments were carried out with protein samples of a strain expressing the fusion protein SltA::HA_3_ (Figure 1B; [20]). As expected, the three main forms of SltA were immunodetected in mycelia grown under standard culture conditions (AMM): the full-length form of 78 kDa, SltA^78kDa^, and a proteolysed version of 32kDa, SltA^32kDa^, also found in a phosphorylated state, SltA^32kDa^-P (C: control, in Figure 1B; [20]). Addition of 1.0 M sodium (as NaCl) or medium alkalinisation (pH 8) by addition of 100 mM Na_2_HPO_4_ caused differential effects on the relative quantities, and probably on the total amount, of these SltA forms. Presence of 1.0 M Na^+^ in the medium induced a reduction in the levels of SltA^78kDa^ along the first 30 min, followed by an increase after 120 min. In contrast, medium alkalinisation caused a transient increase in the levels of SltA^78kDa^ at 30 min. Both stresses induced at 30 min a peak of SltA^32kDa^-P, the proteolysed and phosphorylated form (Figure 1B). These results indicated that adaptation to high sodium concentrations or pH 8 may require differential responses by the different forms of the transcription factor SltA within 1 hour after induction of stress. To investigate the effects of these stresses and the role of SltA on the pattern of gene expression, we decided to extract and sequence RNA samples after 60 minutes of induction of sodium or alkaline pH stress.

### Transcriptional profiles under alkaline pH or sodium stress

Since growth of the *sltA*Δ strain under sodium or alkaline pH stress was completely inhibited on solid medium (Figure 1A), sample collection was carried out from liquid cultures after addition of the compound causing stress (Figure 2A; see also Methods). Mycelial samples cultured in AMM were used as controls. Importantly, mycelial cultures had a pH value of approximately 4 before the addition of sodium or pH 8 shift. Both control and sodium-stress induced cultures maintained the same acidic pH at the time of sample collection. RNA was extracted in biological duplicates and sequenced (see Methods). PCA and sample distance plots showed that sample duplicates clustered together, with a higher similarity between sodium stress and control samples than those corresponding to alkaline pH stress (Figure S1A and S1B). The correlation values obtained followed the general guidelines for biological duplicates, since in all cases they were between 0.94 and 0.97 (higher than 0.90; https://www.encodeproject.org/).

Five transcriptional comparisons were carried out with expression data (Figure 2B; see also the dispersion of log fold change values in the MA-plots of Figure S1C). First, the transcriptomes of the reference wild-type strain grown in alkalinised and in standard AMM media were compared (number 1 in Figure 2B). The expression of 1,248 genes showed a log2FC value higher than 2 or lower than −2, and were considered as significantly de-regulated. That means nearly 10% of the genes annotated in the 36^th^ version of the *A. nidulans* genome. Of these, 387 genes were found upregulated (log2FC > 2) and 861 genes downregulated (log2FC < −2) at pH 8. The second analysis compared the transcriptome of the wild-type strain grown in AMM supplemented with 1.0 M sodium against the control condition (number 2 in Figure 2B), finding 183 genes up- and 217 genes downregulated. The third set of comparisons used the transcriptomes of the null *sltA* and the reference wild-type strains in the three culture conditions tested (numbers 3: AMM, 4: pH 8; and 5: 1.0 M sodium, in Figure 2B), rendering 348, 750 and 341 upregulated genes, and 764, 735 and 754 downregulated genes, respectively. The number of DEGs (differentially expressed genes) in each transcriptional comparison established the framework for further analyses.

Finally, and considering the affinity of SltA for the target sequence 5’-AGGCA-3’ [5], cumulative-distribution fraction plots were drawn with the aim of correlating in each of the above-described transcriptomic comparisons log2FC values with the number of SltA target sites in gene promoters (groups of promoters with 0, 1, 2, 3, 4 and >5 SltA target motifs in the first 1,000 bp upstream of the starting ATG were differentiated) (Figure 2C). General results suggest, on the one hand, that mainly the response to pH 8 but also that to NaCl do not exclusively depend on the number of putative SltA binding sites on target promoters (first two plots in Figure 2C). In all probability, this occurs because SltA-independent regulatory mechanisms are also required for alkaline pH and sodium homeostasis. And on the other hand, that an increase in the number of SltA target sequences in a given promoter correlates with a stronger dependency on the positive regulatory activity of SltA (lower log2FC values) in any of the three culture conditions tested (*sltA*Δ vs WT comparisons in Figure 2C).

### Alkaline pH causes a deregulation of genes coding for membrane transporters, with different levels of dependence on SltA activity

In the wild-type strain, the number of downregulated genes at pH 8 was twice that of upregulated genes (861 vs 387 genes), suggesting that adaptation to medium alkalinisation after shift from an acid culture involves induction but mainly repression of specific sets of genes. Analyses of the lists of significantly up- and downregulated genes (Supplementary Table S3; see also a specific bioinformatic analysis of genes with the Top100 log2FC values in Supplementary Tables S4 and S5) confirmed the previously described upregulation of “alkaline-expressed genes” such as *pacC, aflR* and *enaA* [2, 5, 38, 39], and the downregulation of “acid-expressed genes” such as *palF* [40] (see below). These results support the reliability of the expression values obtained by RNA sequencing. Nevertheless, the expression of *sltA/An2919* itself was high and was not significantly altered in the wild-type strain, suggesting that the transcription factor plays important roles in the three culture conditions studied.

The heatmap in Figure 3A includes all the genes up- and downregulated at pH 8 in the reference wild-type strain but was built considering their row Z scores in all conditions and genetic backgrounds tested. The upper clade in the tree contains genes that are upregulated and the lower clade those downregulated at pH 8 (blue and red colors, respectively). Multiple gene expression patterns could be differentiated. Highlighted are five main expression clusters among upregulated genes and three among downregulated genes at alkaline pH. Those clusters of gene expression also uncovered different dependency levels on SltA activity. Clusters B to D (Figure 3B-3D) correspond to those genes upregulated in the wild-type strain at pH 8. We differentiated three patterns of expression upon SltA activity: i) those in panel B being their transcription partially dependent on SltA (Figure 3B), ii) those in panel C totally dependent on SltA (Figure 3C), and iii) those apparently independent of SltA activity (Figure 3D). These results suggest that the response to alkaline pH implies SltA-dependent and SltA-independent mechanisms and, in some cases, most probably the participation of additional regulators in cooperation with SltA.

**Figure 3:**
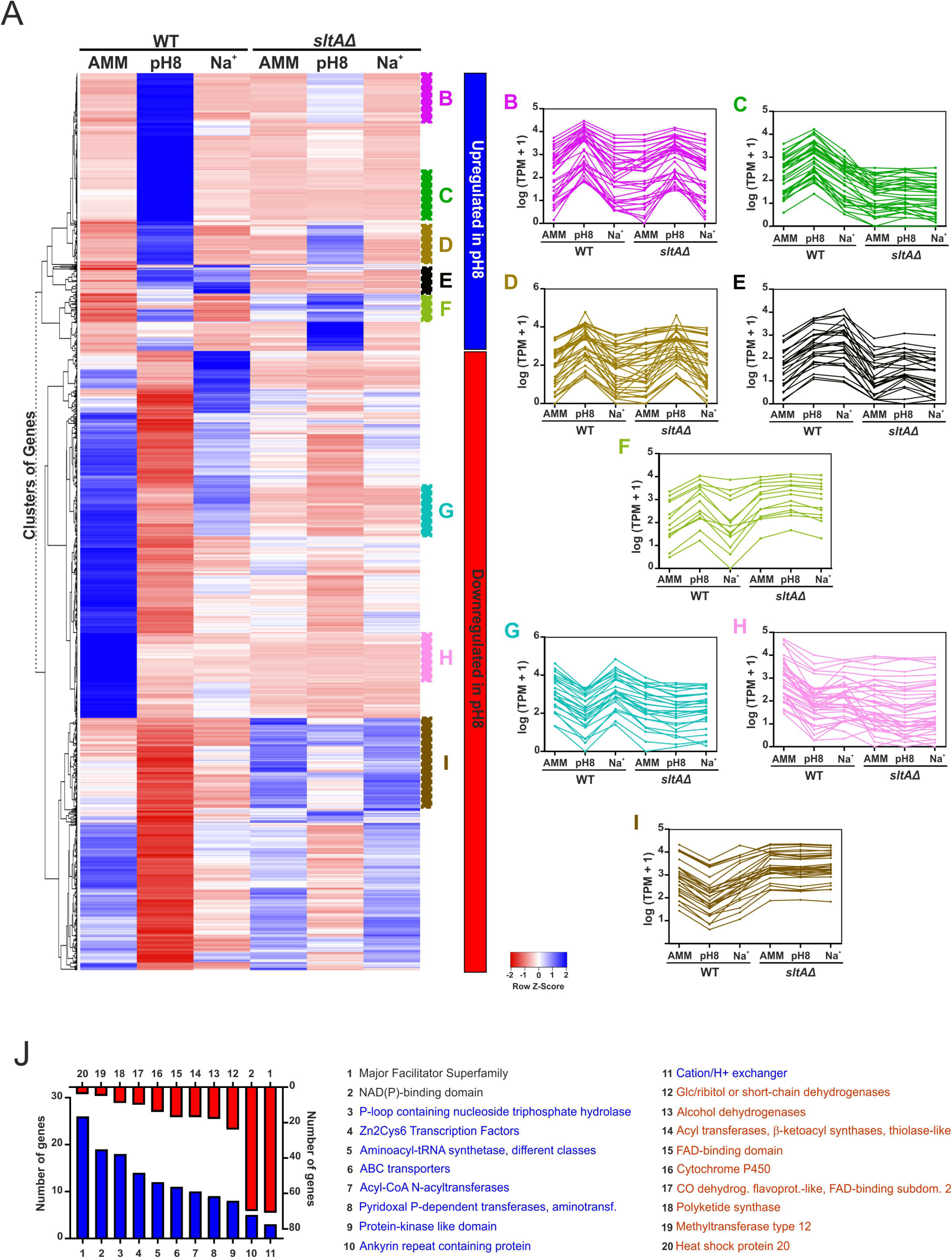
Transcriptional response of *A. nidulans* to alkaline pH. A) Heatmap showing the changes in expression of the genes significantly upregulated (blue color; upper clade of the tree) and downregulated (red color; bottom clade) at pH 8, in all the six conditions and backgrounds tested. B-I) Multiple expression clusters can be differentiated. B: Genes upregulated at pH 8 and partially dependent on SltA at pH 8. C: Genes upregulated at pH 8 and downregulated in the *sltA*Δ background in all conditions tested. D: Genes upregulated at pH 8, almost independently of SltA at pH 8. E: Genes upregulated both at pH 8 and in medium supplemented with 1M Na^+^, SltA-dependent. F: Genes upregulated at pH 8 and upregulated also in the *sltA*Δ background in all conditions tested. G: Genes downregulated at pH 8 but not in culture medium supplemented with 1.0 M Na^+^, and in a SltA-dependent manner. H: Genes downregulated at pH 8 and in the *sltA*Δ background in all conditions tested. I: Genes downregulated at pH 8 but upregulated in the *sltA*Δ background in all conditions tested. J) Bar-plot showing the number of DEGs coding for proteins predicted to contain significantly enriched Interpro domains. Blue color is for significantly upregulated genes at pH 8 while downregulated genes are highlighted in red. Genes coding for Major Facilitator Superfamily transporters are the Top1 up- and downregulated gene family.

Cluster E (Figure 3E) includes genes that are upregulated both at pH 8 and in medium supplemented with 1.0 M Na^+^. This result showed that the responses to these two types of abiotic stress may partially overlap. The expression of genes in Cluster E is SltA-dependent in all conditions tested. To extend the diversity of gene expression patterns at pH 8, we show a small cluster (Figure 3F) in which most genes are upregulated at pH 8 in the wild-type strain and also in the *sltA*Δ strain in all conditions tested.

Among downregulated genes at pH 8, we point to three interesting clusters. Those genes whose expression is downregulated at pH 8 but not under 1.0 M Na^+^ stress (Cluster G in Figure 3G), and those genes downregulated in both stress conditions (Cluster H in Figure 3H). Notably, SltA activity is required for expression of those genes in any condition. The third cluster shows an opposed effect of lacking SltA activity, since in the wild-type strain these genes are downregulated at pH 8 but upregulated in the *sltA*Δ background in all conditions tested (Cluster I in Figure 3I). The overall data suggest that besides acting as an inducer, SltA may have a role as a repressor of transcription.

Finally, we analyzed the enrichment of Interpro domains within proteins coded by genes significantly up- or downregulated at pH 8 (Figure 3J). In both groups, Major Facilitator Superfamily (MFS) and NAD(P)-binding domain-containing proteins were the Top1 and Top2 families, suggesting that the response to pH 8 implies a significant reorganization of the plasma membrane (and other subcellular membranes) and its transporters. In this sense, of note is also the enrichment of ABC transporters and Cation/H^+^ exchangers among proteins coded by genes upregulated at pH 8. Other enriched Interpro domains were, for example, those related to primary and secondary metabolism among the downregulated set of genes and transcription factors (mainly Zn cluster-type TFs) among the upregulated set.

### Transcriptional response to sodium stress

A similar approach was followed to analyze RNA sequencing data of the samples corresponding to sodium stress compared to the control condition. In this case, the number of significantly up- or downregulated genes is clearly lower than the figures described for pH 8, suggesting that sodium stress has a lesser impact on the transcriptome of *A. nidulans*. Similarly, a preliminary analysis of the lists of significantly up- and downregulated genes (Supplementary Table S6; see also a specific bioinformatic analysis of Top100 genes in Supplementary Tables S7 and S8) showed that among previously known SltA-dependent and/or stress-response genes, only *sB*, coding the principal sulfate transporter, varied (upregulation) significantly its expression under sodium stress (see, for example, the upregulation of *pacC, enaA* or *aflR*, but not that of *sB*, at pH 8).

Six gene expression clusters are highlighted in the heatmap of Figure 4A (see panels B-G), two of them corresponding to the clade of upregulated and the remaining four to the clade of downregulated genes. Among upregulated genes, we identified two clusters dependent on SltA activity and also activated by sodium stress. Cluster B shows the presence of multiple genes that are upregulated mainly or exclusively under sodium stress (Figure 4B). And Cluster C highlights genes that are upregulated both at pH 8 and in 1.0 M Na^+^ stress (Figure 4C). Those genes would be common to both response mechanisms (see also Figure 3E).

**Figure 4:**
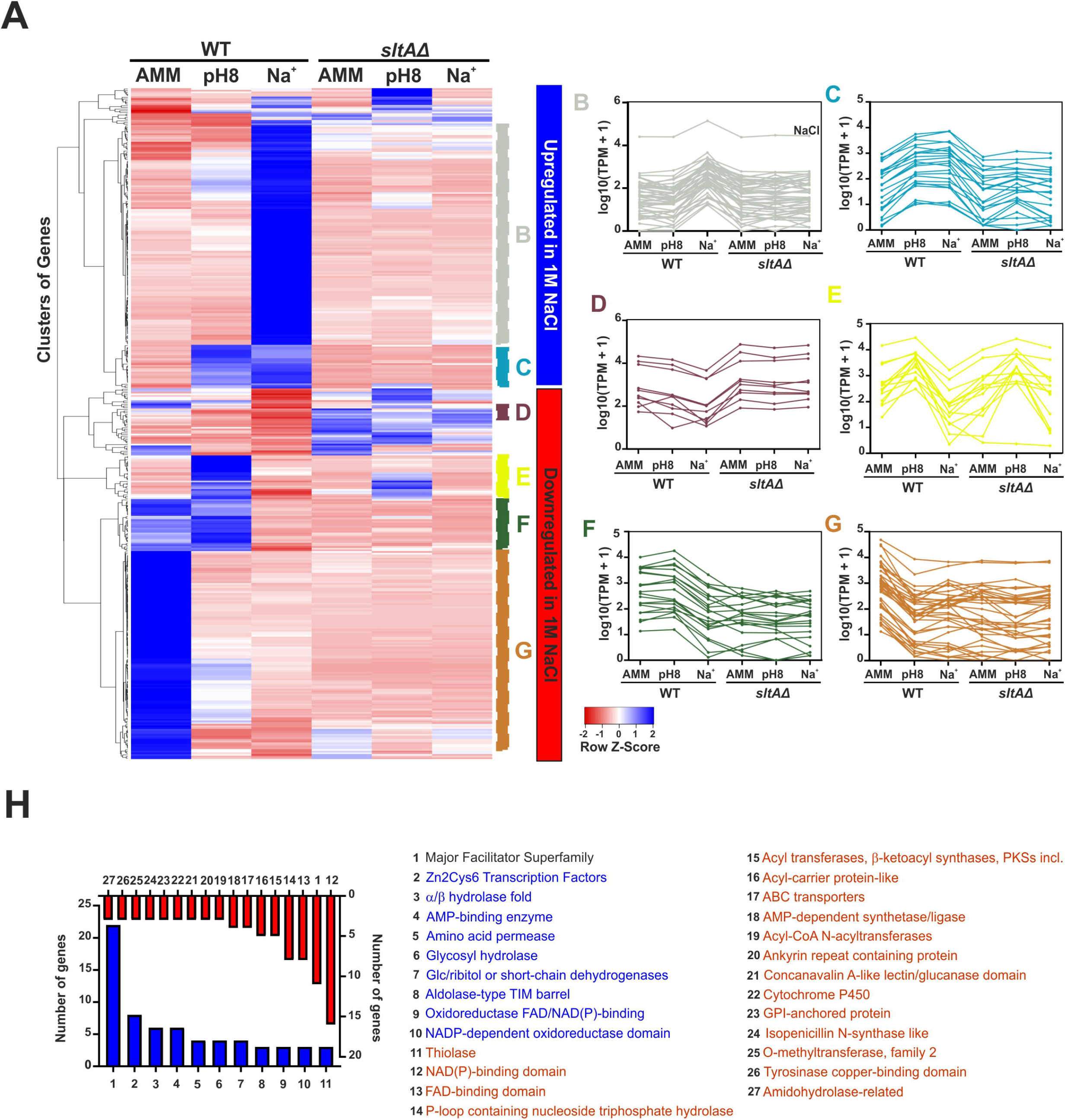
Transcriptional response of *A. nidulans* to sodium (1.0 M NaCl) stress. A) Heatmap showing the changes in expression of the genes significantly upregulated (blue; upper clade) and downregulated (red; bottom clade) in medium supplemented with 1.0 M Na^+^, in all the six conditions and backgrounds tested. B-I) Again, multiple expression clusters can be differentiated. B: Genes upregulated mainly or exclusively under sodium stress and in a SltA-dependent way. C: Genes upregulated both at pH 8 and in medium supplemented with 1.0 M Na^+^, SltA-dependent. D: Genes downregulated at pH 8 but mainly in 1.0 M Na^+^ but upregulated in the *sltA*Δ background in all conditions tested. E: Genes upregulated at pH 8 but downregulated in medium supplemented with 1.0 M Na^+^ (different dependency patterns on SltA at pH 8). F: Genes downregulated only in medium supplemented with 1.0 M Na^+^, SltA-dependent in standard minimal medium and at pH 8. G: Genes expressed in AMM and downregulated in the rest of conditions and backgrounds tested. J) Bar-plot showing the number of genes coding for proteins predicted to contain significantly enriched Interpro domains. Blue color is for significantly upregulated genes under Na^+^ stress while downregulated genes are highlighted in red. As at pH 8, genes coding for Major Facilitator Superfamily transporters are the Top1 up- and downregulated gene family.

Among downregulated genes, cluster D contains genes that are downregulated at pH 8 but mainly in 1.0 M Na^+^, however were upregulated in the *sltA*Δ background in all conditions tested (Figure 4D). Cluster E groups genes that are upregulated at pH 8 but downregulated in medium supplemented with 1.0 M Na^+^, with different dependency patterns on SltA activity at pH 8 (independent or with positive dependence of SltA; Figure 4E). Cluster F designates genes downregulated only in medium supplemented with 1.0 M Na^+^ and that are strongly dependent on the positive SltA activity in AMM and at pH 8 (Figure 4F). Finally, genes in cluster G are expressed in AMM and downregulated in the rest of conditions and backgrounds tested (Figure 4G).

The significantly enriched Interpro domains predicted in the proteins encoded by genes with up- or downregulation under sodium stress (Figure 4H), differed from those described in the case of pH 8 (see Figure 3J for a comparison). However, other families such as MFS (Top1, upregulated), NAD(P)-binding domain or Zn binuclear cluster-type TFs (Top2, upregulated) were common to the response to alkaline pH. Thus, we decided to analyse more deeply the sets and specific subsets of genes significantly deregulated at pH 8 and under sodium stress.

### Transcriptional responses under alkaline pH and sodium stress differ significantly

Only 43 genes were upregulated in both comparisons from a total number of 387 (pH 8) and 183 (1.0 M Na^+^), respectively (only 3 when Top100 tables were compared; Figure 5A). Those genes upregulated simultaneously included the ethanol regulon [41], which extends from An8977 to An8982 in ChrVII (only An8982 was not significantly upregulated in both conditions). This observation, as well as the upregulation of genes coding for MFS proteins annotated as solute:H^+^ symporters having a role in glycerol transport (such as *An9168*, the ortholog of *slt1* of *Saccharomyces cerevisiae*, which is induced by osmotic shock [42]), may be related to an osmoregulation mechanism induced by pH 8 and mainly sodium stress. Indeed, it has been described in the yeast *Pichia sorbitophila* that the ethanol regulon is induced by glycerol accumulated in response to sodium stress and that the main role of this system is osmoregulation, by accumulation of glycerol inside cells to compensate stress caused by alkaline pH or NaCl [43].

**Figure 5:**
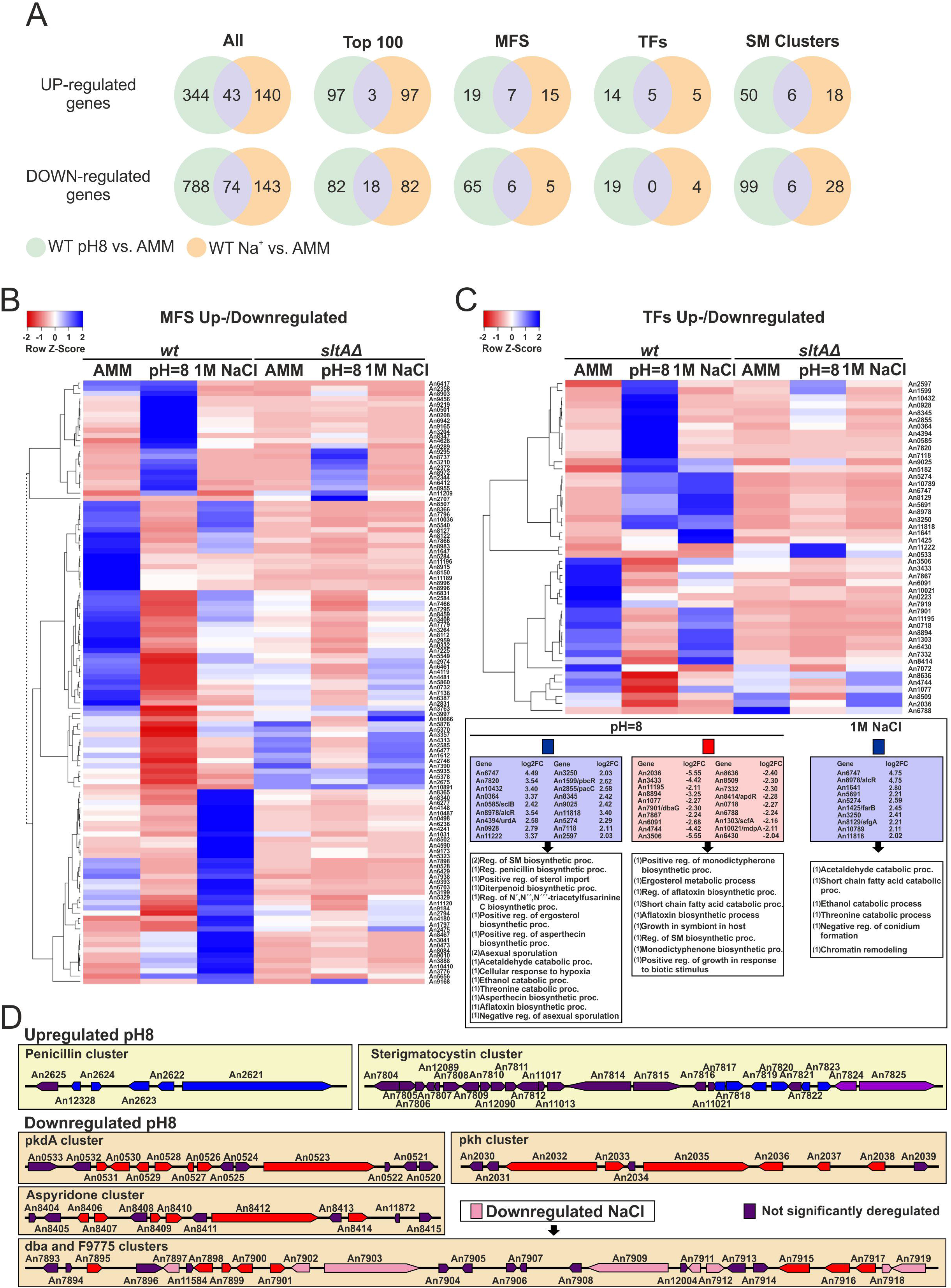
Differential transcriptional responses of *A. nidulans* to alkaline pH or cation stress. A) Venn diagrams showing specific and common significantly deregulated genes when comparing the effect of medium alkalinisation and high sodium stress. “All” indicates the complete lists of DEGs. “Top100” indicates analyses restricted to the first 100 DEGs of each comparison. The rest of Venn diagrams focus on the up-/downregulation of genes coding for Major Facilitator Superfamily proteins (MFS), Transcription Factors (TF) and Secondary Metabolite gene Clusters (SM Clusters). B-C) Heatmaps showing the changes in expression of the genes that, coding for MFS transporters (B) or TFs (C), are significantly deregulated at pH 8 and/or in 1.0 M Na^+^, in all the six conditions and backgrounds tested. Results suggest a very low degree of overlap between both responses regarding genes coding for these two types of proteins. D) Diagrams showing SM Clusters significantly deregulated exclusively at pH 8 or both at pH 8 and in 1.0 M Na^+^.

The number of simultaneously deregulated genes at pH 8 and 1.0 M Na^+^ increased to 74 in the sets of downregulated genes, from a total number of 861 (pH 8) and 217 (1.0 M Na^+^), respectively (only 18 when Top100 tables were compared; Figure 5A). Again, we found little overlap when up- and downregulated genes coding for MFS or TFs were compared (Figure 5A) or in the heatmaps shown in Figure 5B and 5C (see for example genes coding for TFs that are upregulated both at pH 8 and 1.0 M Na^+^ in a SltA-dependent manner, such as *An8978/alcR*, coding for the TF of the ethanol regulon, *An11818* or *An6747*). The differences in the transcriptional responses to alkaline pH and sodium stress were also supported by the analysis of those genes coding for proteins involved in secondary metabolite (SM) production (Figure 5A). Panel D shows the deregulation of mainly different subsets of SM genes, such as the upregulation of penicillin and sterigmatocystin cluster genes at pH 8 [39, 44], or the downregulation of *pkdA*, aspyridone and *pkh* clusters also at pH 8, with only the *dba* cluster being downregulated also under sodium stress [45–48]. Overall, results strongly suggest that the transcriptional responses to sodium and alkaline pH stress differ significantly.

### Deletion of *sltA* causes a profound reprogramming of gene expression, with specific subsets of DEGs depending on culture conditions

Next, we focused on the effect of *sltA* deletion on gene expression, based on three transcriptomic comparisons with the reference wild-type strain: under standard culture conditions (AMM), at pH 8 and after addition of 1.0 M NaCl. In this case, we found that 139 genes were significantly upregulated and 295 downregulated simultaneously in the three transcriptomic comparisons, with additional subsets of genes being deregulated in two or only one of the comparisons (Figure 6A and 6B; see also Venn diagrams corresponding to Top100 lists of DEGs). These figures suggest that deletion of *sltA* has a general effect on gene expression independently of culture conditions but also that there are most probably specific roles in transcriptional regulation at alkaline pH or sodium stress, as suggested before.

**Figure 6:**
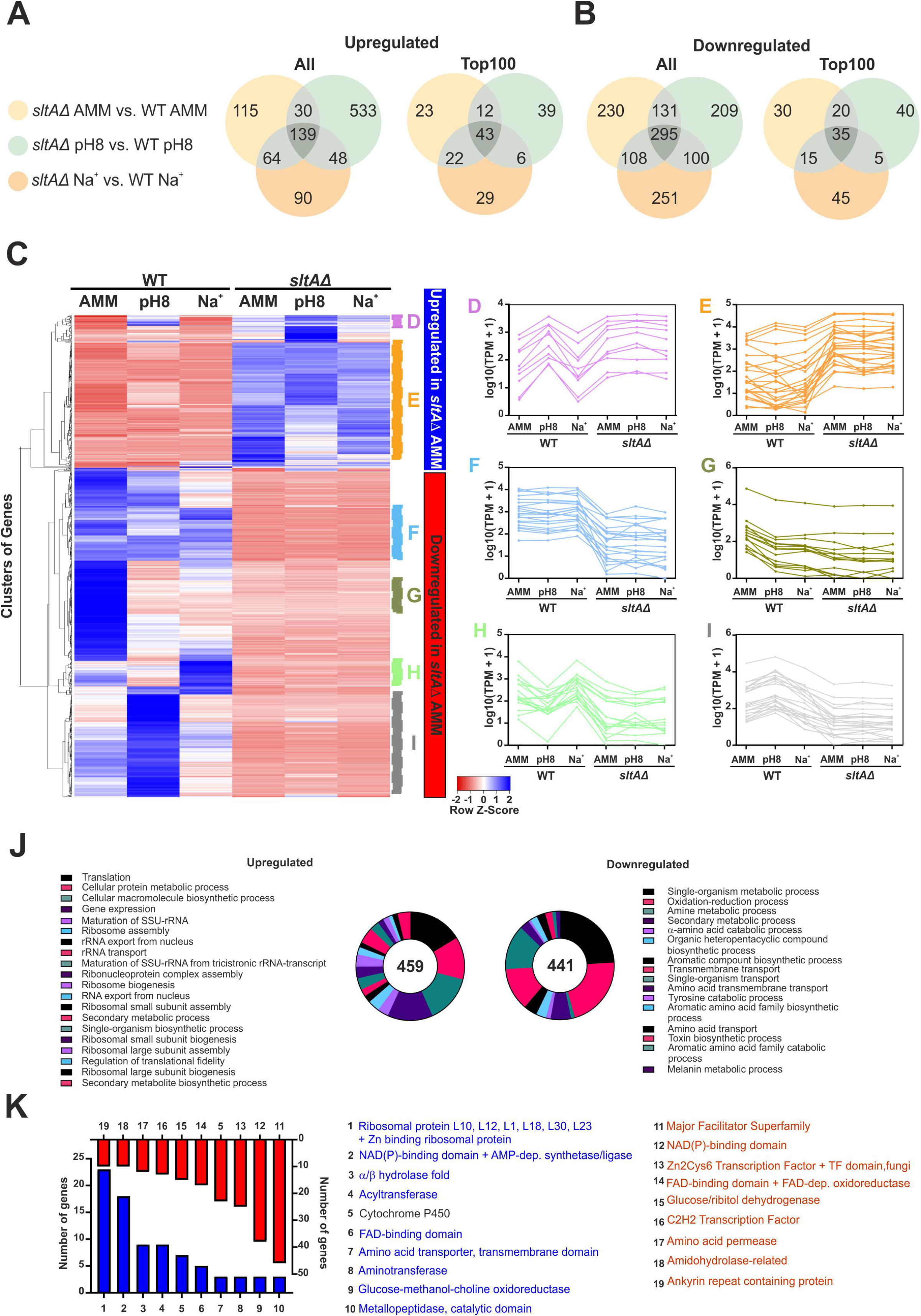
Comparison of *sltA*Δ and wild-type transcriptomes under standard culture conditions. A-B) Venn diagrams showing the number of genes significantly upregulated (A) or downregulated (B) when comparing the *sltA*Δ strain and the reference wild type in any of the three culture conditions analyzed: AMM, AMM medium alkalinised to pH 8, or AMM supplemented with 1.0 M NaCl. Left: Diagram corresponding to the complete lists of DEGs. Right: Diagram corresponding to the Top100 DEGs. C) Heatmap showing the changes in expression of the genes significantly upregulated (blue color; upper clade) and downregulated (red color; bottom clade) in the null *sltA* strain in AMM. Expression in all the six conditions and backgrounds tested is shown. D-I) Multiple expression clusters are differentiated. D: Genes upregulated in all conditions in the *sltA*Δ strain and at pH 8 in the reference wild-type strain. E: Genes with lower expression levels in the wild-type strain compared to the null *sltA* mutant, in all conditions. F: Genes downregulated in the *sltA* mutant in all conditions tested. G: Genes with higher expression levels in the wild-type strain in AMM but downregulated in the rest of samples. H: Genes downregulated in all conditions in the *sltA*Δ strain and at pH 8 in the reference wild-type strain. I: Genes upregulated at pH 8 in the wild-type strain but downregulated in the null *sltA* background in all conditions. J) GO enrichment analyses of the genes up- (left) or downregulated (right) when comparing wild-type and Δ*sltA* samples in AMM. K) Bar-plot showing the number of DEGs coding for proteins predicted to contain significantly enriched Interpro domains. Blue color is for significantly upregulated genes in the *sltA*Δ strain in AMM while downregulated genes are highlighted in red.

Again, multiple expression clusters could be differentiated from the expression patterns of significantly deregulated genes when *sltA*Δ and wild-type samples cultured in AMM were compared (Figure 6C-6I). In the tables of genes corresponding to this comparison (Supplementary Table S9; see also a specific bioinformatic analysis of Top100 genes in Supplementary Tables S10 and S11), we found that previously known SltA-related genes such as *enaA, aflR* or *brlA* were significantly downregulated and putative vacuolar calcium ATPases *pmcA* and *pmcB* were upregulated in the null *sltA* strain [5, 49, 50]. Cluster D corresponds to genes upregulated in all conditions in the *sltA*Δ strain and at pH 8 in the reference wild-type strain (Figure 6D). Cluster E highlights genes with lower expression levels in the wild-type strain compared to the null *sltA* mutant (Figure 6E), in all conditions, and probably corresponds to the subset of 139 genes mentioned in Figure 6A (left). The remaining four expression clusters highlighted in the heatmap of Figure 6 correspond to genes that are downregulated in the null *sltA* strain in AMM (lower clade of the heatmap). Cluster F shows genes that are downregulated in the *sltA* mutant in all conditions tested (Figure 6F). Cluster G includes those genes with higher expression levels in the wild-type strain in AMM but downregulated in the rest of samples (Figure 6G). Cluster H highlights those genes downregulated in all conditions in the *sltA*Δ strain and at pH 8 in the reference wild-type strain (Figure 6H). And finally, Cluster I groups genes upregulated at pH 8 in the wild-type strain but downregulated in the null *sltA* background in all conditions (Figure 6I).

GO enrichment analyses showed that most GO terms among upregulated genes were related to ribosome biogenesis and translation (Figure 6J), suggesting that deletion of *sltA* caused a ribosomal stress that would correlate with the growth and developmental phenotype of the null strain on AMM (see Figure 1A). Also, we found a majority of GO terms related to primary and mainly secondary metabolism among deregulated genes. A hypothetic deregulation of secondary metabolism would correlate with an induction of ribosomal stress [47, 48]. These results were confirmed by the Interpro domains enriched in the proteins coded by DEGs (Figure 6K). Of note is, again, the downregulation of MFS transporter-coding genes, those coding for NAD(P)-binding proteins and those coding for TFs, suggesting that the previously mentioned deregulation of these groups of genes depends, at least partially, on SltA.

In the tables of genes corresponding to the comparison between *sltA*Δ and wild-type strains at pH 8 (Supplementary Table S12; see also a specific bioinformatic analysis of Top100 genes in Supplementary Tables S13 and S14) or sodium stress (Supplementary Tables S15, S16 and S17), we found that previously known SltA-regulated genes such as *vcxA, pmcA* and *pmcB* were significantly upregulated while others such as *sltB, aflR, enaA* and *brlA* were downregulated [5, 21, 49, 50]. The clusters inferred from the heatmaps in supplementary Figures S2A and S3A confirmed most of the gene expression patterns described above for the comparison of the same strains in AMM (see Supplementary Figures S2B-S2H and S3B-S3J). In this case, and as described for the wild-type strain, genes coding for MFS transporters, NAD(P)-binding domain proteins and TFs (Zn2Cys6 and C2H2 types) were among the most significantly deregulated ones (Supplementary Figures S2H and S3K). Overall, results suggest that SltA has a broad positive role in the control of genetic pathways.

### SltA function is required for proper expression of *pacC* and *palF* genes

RNA sequencing data agreed with previous results that showed *pacC* as an alkaline-expressed gene and *palF* as an acid-expressed gene [2, 40]. In this opposed regulation PacC has been suggested to play an important role in an auto-regulatory model of the ambient pH regulatory system. However, expression levels of *pacC* and *palF* were markedly altered in the null *sltA* background. RNAseq data evidenced that up-regulation of *pacC* expression caused by medium alkalinisation was lost in the *sltA* mutant (Figure 7A) without being affected those levels found at acid pH (AMM). In contrast, *palF* expression was notably reduced at acid pH, close to those levels found at alkaline pH (Figure 7B). To verify this finding, independent biological repetitions were done and *pacC* and *palF* expression was determined by means of quantitative PCR. Results of qPCRs, using α-tubulin coding gene *benA* as control, were similar to those found in RNA sequencing experiments (Figure 7, compare RNA-seq and qPCR charts). Absence of SltA activity caused deregulation of these principal genes in the ambient pH regulatory system.

**Figure 7:**
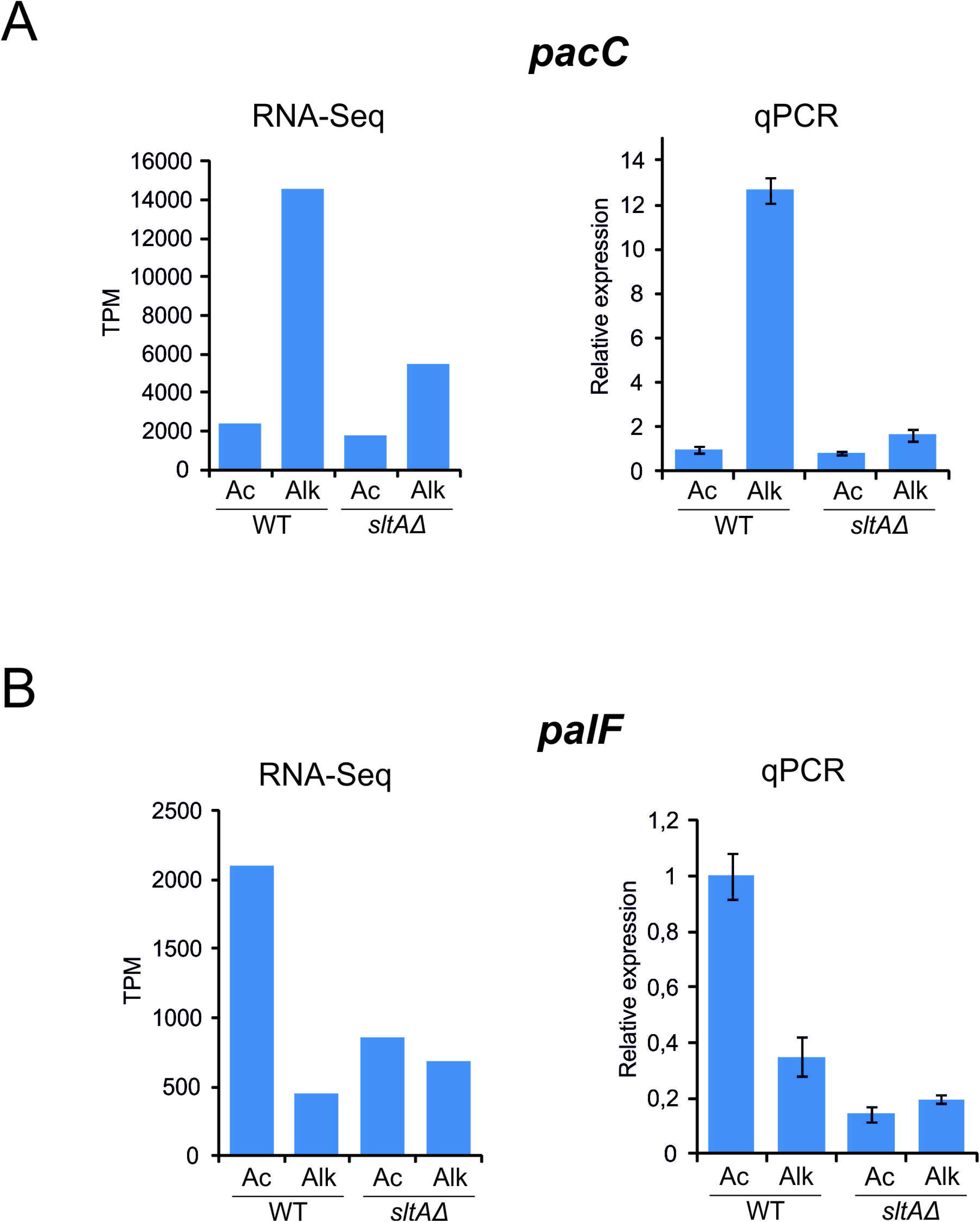
Analyses of *pacC* and *palF* expression levels under alkaline pH stress and the effect of lacking SltA activity. Confirmation of RNA sequencing data in new biological samples using quantitative PCR. A) Charts showing RNA sequencing data (TPM values) and relative expression levels of *pacC* in wild-type and *sltA*Δ strains, in control AMM (Ac) and pH 8 (Alk) conditions. B) As in A) but for *palF* gene expression levels. Strains used in qPCR experiments were MAD6669 (reference wild-type strain) and MAD3816 (*sltA*Δ).

## Discussion

Tolerance to abiotic stress conditions has served as a basis for classification of microorganisms. The capacity to grow at a given pH results in differentiating acidophiles, neutrophiles and alkaliphiles, while those organisms able to thrive when alkali cations, such as sodium, are present in high quantities are known as halo-tolerants (recently reviewed in [51]). Of course, tolerance relies on genomes coding for subsets of proteins enabling adaptation to those abiotic stress conditions and also the induction of regulatory mechanisms that coordinate the scheduled expression of those resistance genes [52]. Undoubtedly, *A. nidulans* has served as an important reference system to investigate these abiotic stress-responsive pathways (recently reviewed in [1]).

In this work, we have focused on studying the transcriptional response to alkalinisation of the extracellular medium and to high concentrations of extracellular sodium. Both stress conditions threaten the colonial growth capacity of *A. nidulans*, particularly when the function of the transcriptional factor SltA is lost (Figure 1A; [5, 20]). Here we show that *A. nidulans* modulates differently over time the relative amounts of the three main forms of SltA, SltA^78kDa^ as well as phosphorylated and non-phosphorylated SltA^32kDa^ forms, after the induction of each stress condition (Figure 1B). This observation suggests that all forms of SltA may play a role in controlling gene expression, and that specific relative amounts are required to coordinate the adequate transcriptional response to each stimulus. To determine this transcriptional response, we concentrated at the time point of 60 minutes after induction of stress because major differences in the kinetics of SltA 78kDa and 32kDa forms were observed at 30 minutes (Figure 1B).

Comparisons of RNA sequencing results in the wild type strain have revealed important transcriptomic differences between the responses to medium alkalinisation and sodium stress. The PCA plot (Supplementary Figure S1) denotes that ambient pH alkalinisation causes a major modification of the transcriptional pattern, meanwhile addition of sodium has a more specific effect. Subsequent comparisons made support the above conclusion, with the observation that only a few genes are simultaneously deregulated in the wild type strain when each of the two types of stress is induced (Figure 5A). Among upregulated genes, 43 are common but 344 are specific of alkaline pH response and 140 of high sodium response. On the contrary, 74 genes are downregulated in both conditions and 788 and 143 genes are specifically downregulated at pH8 and high sodium, respectively. Part of this common response are the ethanol regulon and glycerol/H^+^ symporters. Accumulation of osmolytes such as glycerol or trehalose is a common mechanism of osmoregulation [53]. Although the anabolic pathway of glycerol (or the expression of *vosA*, which codes for a transcriptional regulator of trehalose synthesis [54]) is not found in the list of DEGs, the upregulation of the ethanol regulon and glycerol/H+ symporters suggests that intracellular accumulation of glycerol is probably a primary response to these abiotic stresses in *A. nidulans*, more prominent in the case of high sodium concentrations.

Intracellular production of glycerol as an osmolyte is regulated by the Mitogen activated protein kinase cascade HOG, HogA/SakA in *A. nidulans*, through the transcription factor MsnA [38, 55, 56]. In the wild type strain, none of the genes encoding elements of this pathway (reviewed in [52, 57, 58]) are included in the DEG lists of this work. This is interesting because the absence of any of the elements of the MAP kinase cascade, *sskB* (MAPKKK), *pbsB* (MAPKK) and *hogA* (MAPK) cause sensitivity to high NaCl [57]. Our results indicate that this key signaling pathway in osmoregulation is not subjected to transcriptional regulation during early stages of the response to alkaline or sodium stress. Then, which is the main transcriptional response observed in the wild type? At alkaline pH, among the most upregulated genes are those coding for sodium ATPases, *enaA* (AN6642) and *enaB* (AN1628), the Pi/Na^+^ symporter Pho89-like (AN8956), and two sodium/H^+^ antiporters (AN5035 and AN4131) of the Nha1 family [5, 38, 59]. The upregulation of these ion transporters could reflect the need for sodium and phosphate detoxification or for the accumulation of protons to compensate in intracellular alkalinisation caused by the addition of the buffer. However, this response is specific of alkaline pH because these genes are not significantly deregulated in the response to sodium. In contrast, sodium generates a more general response: the previously described activation of ethanol regulon and a deregulation of genes related to primary metabolism of carbon and nitrogen sources. This effect on primary metabolisms has been previously described as a general response to environmental stress [52, 58].

Which is the role of SltA in transcription? The absence of SltA function causes evident changes in the pattern of gene expression compared to the wild type strain, both in the absence or presence of alkaline pH/sodium stress conditions. This is well observed in the PCA plot in Supplementary Figure S1A, in which data from the null strain do not cluster with and are distant from those of the reference strain. The need for SltA activity under non-stressing conditions is well supported by the fact that the three forms of SltA are detected (Figure 1B). However, the same experiment suggests a requirement of the full-length SltA form at early time-points after alkaline pH stress induction, while the truncated 32kDa form predominates at high sodium concentrations. These observations might explain previous results in which the sole expression of a form mimicking the SltA^32kDa^ truncated version suffices for enabling wildtype growth rates in sodium stress, but not at alkaline pH [20]. Thus, SltA^73kDa^ may play a role in the response to alkalinity and sltA^32kDa^ under both stress conditions.

More than a thousand of genes are likely to be significantly deregulated in each of the three wild-type versus *sltA*Δ comparisons (i.e. 1,112 genes under standard, non-stressing, culture conditions; Figure 2B) and may require the direct or indirect activity of SltA. Among them, we found genes that are up- and downregulated, confirming our previous hypothesis that SltA may play a dual role as a repressor and as an activator [5, 49]. RNA sequencing data confirms that SltB, coding for the protease that cleaves SltA, is among the genes positively dependent on SltA activity [20]. Other known targets of SltA are also present among DEGs (see results). However, the most representative GO classes upregulated in the null *sltA* strain under non-stressing conditions are related to ribosome biogenesis and translation (Figure 6J-K), indicating that the absence of SltA forces a mis-scheduled regulation of the expression of these genes. The deregulation of expression of ribosome-forming or regulatory proteins has been related to a general response to ambient stress [52]. Thus, in the case of the lack of SltA activity, responses specifically controlled by SltA and those general to environmental stresses should be differentiated.

Several transcriptional regulators are subjected to SltA control. Expression of *msnA*, a transducer of HOG-osmoregulatory pathway [38], is significantly downregulated in the null *sltA* background. In *S. cerevisiae, ALD2* gene, coding for cytoplasmic aldehyde dehydrogenase [60], is under regulation of Msn1p, the MsnA homologue. In concordance, the homologue of ALD2 in *A. nidulans, AN9034*, is strongly downregulated in the null *sltA* background. Predictably the product of *AN9034* converts acetaldehyde in acetyl-CoA and its downregulation in the null *sltA* strain might cause the intracellular elevation of toxic aldehydes, as has been described in yeast [60], contributing to the reduced viability of the *sltA*Δ mutant under the stress conditions analyzed in this work. Other transcription factor coding genes deregulated in the *sltA* deletion strain belong to well-known SM biosynthetic clusters [61], evidencing regulation by ambient stress and/or in concert with SltA regulation. As previously known, the penicillin cluster is one of this alkaline-pH activated SM pathways [44], but only *acvA* (ACV-peptide synthase) and *ipnA* (isopenicillin N synthase), the genes coding for the first two enzymes, are among the Top100 upregulated genes at alkaline pH (Figure 5D). The penicillin pathway is not under SltA regulation, but certainly SltA plays an important role on the regulation of the aflatoxin pathway [50], probably by positively regulating the expression of the zinc binuclear cluster-coding gene *aflR* (see Supplementary Tables S9 and S12). Other deregulated SM pathways at alkaline pH are the *pkh* cluster, most likely through downregulation of the predicted specific regulator AN2036, the *pkdA* cluster, the aspyridone pathway (probably via downregulating the specific transcription factor *apdR*) or upregulation of the terrequinone cluster, via upregulation of *tdiB* and *tdiC* genes. Several genes of contiguous clusters *dba* and F9775 are deregulated under sodium stress (Figure 5D). Thus, abiotic stresses and SltA are key regulators of several SM pathways [61].

In particular, this investigation opens an avenue for defining the hierarchy among the main transcriptional regulators controlling the response to ambient pH. PacC, the transcription factor regulating ambient pH response, is upregulated at alkaline pH, as has been shown before (Figure 5C and Figure 7A) [2, 40]. The absence of SltA function strongly reduces the upregulation of *pacC* at alkaline pH, and this has been confirmed using an alternative null *sltA* strain and through qPCR analyses (Figure 7A). Transcripcional upregulation of *pacC* has been proposed as a key step in the PacC-Pal ambient pH pathway, and thus, the fact that *pacC* transcription levels remain close to those determined at acid pH in the null *sltA* mutant suggests that there is a significant mismatch in the PacC signalling process. In agreement with this idea, also the acid-expressed gene *palF*, coding for the signaling arrestin-like protein [40], is not properly upregulated at acid pH in the *sltA* mutant. Overall, these data suggest an important role of SltA in modulating signaling in the PacC/Pal pathway in response to alkaline pH. Recently PacX, a negative regulator, was included in this pathway [40]. Here, SltA is proposed to act as a positive regulator of *pacC* and *palF* expression. The presence of three putative SltA binding sites at the *pacC* promoter (first 1,000 bp-s; not shown) suggests a direct role of SltA through direct interaction to those sites. Furthermore, here we have described a correlation between the number of SltA target sites in a promoter and an increase in the fold-change values of wild-type versus *sltA*Δ samples, although this is clearly not the only requirement for a direct transcriptional dependency on SltA activity, since there are multiple genes with multiple SltA sites in their promoters but with log2FC values between −1 and 1. On the other hand, no SltA-binding sites are found in the *palF* promoter, suggesting an indirect function or through recognition of alternative target sites. Future work will focus on delimiting the role of SltA on the ambient pH regulatory pathway and whether corregulation with PacX occurs, since both SltA and PacX are transcription factors specific of Pezizomycotina subphylum [5, 20, 40]. In the context of ancient gene rewiring phenomena [62], both factors may have been recruited for new functions in ancestors of Aspergillus species. Deciphering the roles of SltA in distant Pezizomycotina species would contribute to understand these genetic processes and dissect the mechanisms of abiotic stress response in fungi.

## Conclusions

Our data suggest that alkaline pH and sodium stress responses in *Aspergillus nidulans* include mostly exclusive constituents, with only a small fraction of common elements. Adaptation to alkaline pH apparently requires a more general transcriptional response while it is more specific in the case of sodium stress. Both stress responses include probably a general reorganization of lipid bilayers and their transporters, but mainly through exclusive constituents, suggesting that the study of each response will need specific experimental approaches in the future. At the same time, the transcription factor SltA plays a key role in both responses, with links also with the control of, for example, SM and development. Furthermore, there is a hierarchy of authority between SltA and PacC, with the former controlling the expression of the latter. Future studies will elucidate how this dependency was established in *A. nidulans*.

## Author contribution

E.A.E conceived and supervised the experiments. I.P and E.R. performed experimental work. A.G. carried out trimming and mapping of sequencing reads. Bioinformatic analyses of the sequencing results were done by O.E., I.P. and E.A.E. Original draft manuscript written by O.E., I.P and E.A.E. All authors contributed to the improvement of the text and figures.

## Acknowledgements

To Irene Tomico for sharing strain MAD6669.

## Supplementary material

**<u>Supplementary Figure S1</u>:** Analysis of the reproducibility of the samples analyzed by RNA sequencing. A) PCA analysis of the duplicate samples sequenced by RNA sequencing (see Methods). B) Sample distance plot for the same samples as in panel A. C) MA-plots showing the dispersion of log fold change values in each of the transcriptomic comparisons carried out in this study.

**<u>Supplementary Figure S2</u>: Comparison of *sltA*Δ and wild-type transcriptomes at alkaline pH.** A) Heatmap showing the changes in expression of the genes significantly upregulated (blue color; upper clade) and downregulated (red color; bottom clade) in the null *sltA* strain at pH 8. Expression in all the six conditions and backgrounds tested is shown. B-H) Expression clusters inferred from the heatmap in A. H) Bar-plot showing the number of DEGs coding for proteins predicted to contain significantly enriched Interpro domains. Blue color is for significantly upregulated genes in the *sltA*Δ strain at pH 8 while downregulated genes are highlighted in red. Genes coding for Zn2Cys6 and/or fungal specific transcription factors are the Top1 downregulated gene family.

**<u>Supplementary Figure S3</u>: Comparison of *sltA*Δ and wild-type transcriptomes under 1.0 M Na^+^ stress.** A) Heatmap showing the changes in expression of the genes significantly upregulated (blue color; lower clade of the tree) and downregulated (red color; upper clade) in the null *sltA* strain cultured under Na^+^ stress. Expression in all conditions and backgrounds tested is shown. B-J) Expression clusters inferred from the heatmap in A. K) Bar-plot showing the number of DEGs coding for proteins predicted to contain significantly enriched Interpro domains. Blue color is for significantly upregulated genes in the *sltA*Δ strain under sodium stress while downregulated genes are highlighted in red.

**<u>Supplementary Table S1</u>:** Oligonucleotides used in this study.

**<u>Supplementary Table S2</u>:** Strains used in this study.

**<u>Supplementary Table S3</u>:** Genes significantly de-regulated (log2FC higher or lower than 2) under alkaline pH (pH 8) compared to standard culture conditions (final pH of approximately 4).

**<u>Supplementary Table S4</u>:** Top100 genes significantly upregulated under alkaline pH (pH 8) compared to standard culture conditions.

**<u>Supplementary Table S5</u>:** Top100 genes significantly downregulated under alkaline pH (pH 8) compared to standard culture conditions.

**<u>Supplementary Table S6</u>:** Genes significantly de-regulated under stress conditions induced by the addition of 1.0 M NaCl to the culture medium compared to standard culture conditions.

**<u>Supplementary Table S7</u>:** Top100 genes significantly upregulated under stress conditions induced by the addition of 1.0 M NaCl to the culture medium compared to standard culture conditions.

**<u>Supplementary Table S8</u>:** Top100 genes significantly downregulated under stress conditions induced by the addition of 1.0 M NaCl to the growth medium.

**<u>Supplementary Table S9</u>:** Genes significantly de-regulated in a Δ*sltA* background compared to the wild-type background under standard culture conditions.

**<u>Supplementary Table S10</u>:** Top100 genes significantly upregulated in a Δ*sltA* background compared to the wild-type background under standard culture conditions.

**<u>Supplementary Table S11</u>:** Top100 genes significantly downregulated in a Δ*sltA* background compared to the wild-type background under standard culture conditions.

**<u>Supplementary Table S12</u>:** Genes significantly de-regulated in a Δ*sltA* background compared to the wild-type background under alkaline pH stress conditions.

**<u>Supplementary Table S13</u>:** Top100 genes significantly upregulated in a Δ*sltA* background compared to the wild-type background under alkaline pH stress conditions.

**<u>Supplementary Table S14</u>:** Top100 genes significantly downregulated in a Δ*sltA* background compared to the wild-type background under alkaline pH stress conditions.

**<u>Supplementary Table S15</u>:** Genes significantly de-regulated in a Δ*sltA* background compared to the wild-type background under stress conditions induced by the addition of 1.0 M NaCl.

**<u>Supplementary Table S16</u>:** Top100 genes significantly upregulated in a Δ*sltA* background compared to the wild type under stress conditions induced by the addition of 1.0 M NaCl.

**<u>Supplementary Table S17</u>:** Top100 genes significantly downregulated in a Δ*sltA* background compared to the wild type under stress conditions induced by the addition of 1.0 M NaCl.

## References

1. Etxebeste O, Espeso EA. *Aspergillus nidulans* in the post-genomic era: a top-model filamentous fungus for the study of signaling and homeostasis mechanisms. Int Microbiol 2020;23:5–22.

2. Tilburn J, Sarkar S, Widdick DA, Espeso EA, Orejas M, et al. The *Aspergillus* PacC zinc finger transcription factor mediates regulation of both acid- and alkaline-expressed genes by ambient pH. EMBO J 1995;14:779–790.

3. Arst HN, Peñalva MA. pH regulation in *Aspergillus* and parallels with higher eukaryotic regulatory systems. Trends Genet 2003;19:224–231.

4. Peñalva MA, Tilburn J, Bignell E, Arst HN. Ambient pH gene regulation in fungi: making connections. Trends Microbiol 2008;16:291–300.

5. Spielvogel A, Findon H, Arst HN, Araújo-Bazán L, Hernández-Ortíz P, et al. Two zinc finger transcription factors, CrzA and SltA, are involved in cation homoeostasis and detoxification in *Aspergillus nidulans*. Biochem J;414. Epub ahead of print 2008. DOI: 10.1042/BJ20080344.

6. Orejas M, Espeso EA, Tilburn J, Sarkar S, Arst HN, et al. Activation of the *Aspergillus* PacC transcription factor in response to alkaline ambient pH requires proteolysis of the carboxy-terminal moiety. Genes Dev 1995;9:1622–1632.

7. Espeso EA, Tilburn J, Sánchez-Pulido L, Brown C V, Valencia A, et al. Specific DNA recognition by the *Aspergillus nidulans* three zinc finger transcription factor PacC. J Mol Biol 1997;274:466–480.

8. Espeso EA, Roncal T, Díez E, Rainbow L, Bignell E, et al. On how a transcription factor can avoid its proteolytic activation in the absence of signal transduction. EMBO J 2000;19:719–728.

9. Díez E, Álvaro J, Espeso EA, Rainbow L, Suárez T, et al. Activation of the Aspergillus PacC zinc finger transcription factor requires two proteolytic steps. EMBO J 2002;21:1350–1359.

10. Calcagno-Pizarelli AM, Negrete-Urtasun S, Denison SH, Rudnicka JD, Bussink H-J, et al. Establishment of the ambient pH signaling complex in *Aspergillus nidulans:* PalI assists plasma membrane localization of PalH. Eukaryot Cell 2007;6:2365 LP – 2375.

11. Herranz S, Rodríguez JM, Bussink H-J, Sánchez-Ferrero JC, Arst HN, et al. Arrestin-related proteins mediate pH signaling in fungi. Proc Natl Acad Sci U S A 2005;102:12141 LP – 12146.

12. Vincent O, Rainbow L, Tilburn J, Arst Herbert N. J, Peñalva MA. YPXL/I Is a Protein Interaction Motif Recognized by *Aspergillus* PalA and Its Human Homologue, AIP1/Alix. Mol Cell Biol 2003;23:1647 LP – 1655.

13. Galindo A, Hervás-Aguilar A, Rodríguez-Galán O, Vincent O, Arst HN, et al. PalC, one of two Bro1 domain proteins in the fungal pH signalling pathway, localizes to cortical structures and binds Vps32. Traffic 2007;8:1346–1364.

14. Peñas MM, Hervás-Aguilar A, Múnera-Huertas T, Reoyo E, Peñalva MÁ, et al. Further characterization of the signaling proteolysis step in the Aspergillus nidulans pH signal transduction pathway. Eukaryot Cell 2007;6:960 LP – 970.

15. Hervás-Aguilar A, Rodríguez JM, Tilburn J, Arst HN, Peñalva MA. Evidence for the direct involvement of the proteasome in the proteolytic processing of the Aspergillus nidulans zinc finger transcription factor PacC. J Biol Chem 2007;282:34735–47.

16. Espeso EA, Peñalva MA. Three binding sites for the *Aspergillus nidulans* PacC Zinc-finger Transcription Factor are necessary and sufficient for regulation by ambient pH of the Isopenicillin N Synthase gene promoter. J Biol Chem 1996;271:28825–28830.

17. Espeso EA, Arst HN. On the mechanism by which alkaline pH prevents expression of an Acid-Expressed gene. Mol Cell Biol 2000;20:3355 LP – 3363.

18. Caddick MX, Brownlee AG, Arst HN. Regulation of gene expression by pH of the growth medium in Aspergillus nidulans. Mol Gen Genet 1986;203:346–53.

19. Maccabe AP, Orejas M, Pérez-González JA, Ramón D. Opposite patterns of expression of two *Aspergillus nidulans* xylanase genes with respect to ambient pH. J Bacteriol 1998;180:1331 LP – 1333.

20. Mellado L, Arst HN, Espeso EA. Proteolytic activation of both components of the cation stress-responsive Slt pathway in *Aspergillus nidulans*. Mol Biol Cell 2016;27:2598–612.

21. Mellado L, Calcagno-Pizarelli AM, Lockington RA, Cortese MS, Kelly JM, et al. A second component of the SltA-dependent cation tolerance pathway in *Aspergillus nidulans*. Fungal Genet Biol 2015;82:116–128.

22. Saloheimo A, Aro N, Ilmén M, Penttilä M. Isolation of the ace1 gene encoding a Cys(2)- His(2) transcription factor involved in regulation of activity of the cellulase promoter *cbh1* of *Trichoderma reesei*. J Biol Chem 2000;275:5817–25.

23. O’Neil JD, Bugno M, Stanley MS, Barham-Morris JB, Woodcock NA, et al. Cloning of a novel gene encoding a C2H2 zinc finger protein that alleviates sensitivity to abiotic stresses in *Aspergillus nidulans*. Mycol Res 2002; 106:491–498.

24. Cove DJ. The induction and repression of nitrate reductase in the fungus *Aspergillus nidulans*. Biochim Biophys Acta - Enzymol Biol Oxid 1966;113:51–56.

25. Hernández-Ortiz P, Espeso EA. Phospho-regulation and nucleocytoplasmic trafficking of CrzA in response to calcium and alkaline-pH stress in *Aspergillus nidulans*. Mol Microbiol 2013;89:532–551.

26. Martin M. Cutadapt removes adapter sequences from high-throughput sequencing reads. EMBnet.journal; Vol 17, No 1 Next Gener Seq Data Anal - 1014806/ej171200.

27. Stajich JE, Harris T, Brunk BP, Brestelli J, Fischer S, et al. FungiDB: an integrated functional genomics database for fungi. Nucleic Acids Res 2011;40:D675–D681.

28. Dobin A, Davis CA, Schlesinger F, Drenkow J, Zaleski C, et al. STAR: ultrafast universal RNA-seq aligner. Bioinformatics 2012;29:15–21.

29. Liao Y, Smyth GK, Shi W. featureCounts: an efficient general purpose program for assigning sequence reads to genomic features. Bioinformatics 2013;30:923–930.

30. Li B, Dewey CN. RSEM: accurate transcript quantification from RNA-Seq data with or without a reference genome. BMC Bioinformatics 2011;12:323.

31. Love MI, Huber W, Anders S. Moderated estimation of fold change and dispersion for RNA-seq data with DESeq2. Genome Biol 2014;15:550.

32. R Core Team (2017). R: A language and environment for statistical computing. R Found Stat Comput Vienna, Austria. http://www.r-project.org/ (2017).

33. Mitchell AL, Attwood TK, Babbitt PC, Blum M, Bork P, et al. InterPro in 2019: improving coverage, classification and access to protein sequence annotations. Nucleic Acids Res 2018;gky1100–gky1100.

34. Priebe S, Kreisel C, Horn F, Guthke R, Linde J. FungiFun2: a comprehensive online resource for systematic analysis of gene lists from fungal species. Bioinformatics 2014;31:445–446.

35. Ge SX, Jung D, Yao R. ShinyGO: a graphical gene-set enrichment tool for animals and plants. Bioinformatics. Epub ahead of print 2019. DOI: 10.1093/bioinformatics/btz931.

36. Babicki S, Arndt D, Marcu A, Liang Y, Grant JR, et al. Heatmapper: web-enabled heat mapping for all. Nucleic Acids Res 2016;44:W147–W153.

37. Heberle H, Meirelles GV, da Silva FR, Telles GP, Minghim R. InteractiVenn: a web-based tool for the analysis of sets through Venn diagrams. BMC Bioinformatics 2015;16:169.

38. Han K-H, Prade RA. Osmotic stress-coupled maintenance of polar growth in *Aspergillus nidulans*. Mol Microbiol 2002;43:1065–1078.

39. Delgado-Virgen F, Guzman-de-Peña D. Mechanism of Sterigmatocystin biosynthesis regulation by pH in *Aspergillus nidulans*. Braz J Microbiol 2009;40:933–942.

40. Bussink H-J, Bignell EM, Múnera-Huertas T, Lucena-Agell D, Scazzocchio C, et al. Refining the pH response in *Aspergillus nidulans:* a modulatory triad involving PacX, a novel zinc binuclear cluster protein. Mol Microbiol 2015;98:1051–1072.

41. Felenbok B, Sequeval D, Mathieu M, Sibley S, Gwynne DI, et al. The ethanol regulon in Aspergillus nidulans: characterization and sequence of the positive regulatory gene alcR. Gene 1988;73:385–396.

42. Ferreira C, van Voorst F, Martins A, Neves L, Oliveira R, et al. A member of the sugar transporter family, Stl1p is the glycerol/H+ symporter in Saccharomyces cerevisiae. Mol Biol Cell 2005;16:2068–2076.

43. Lages F, Lucas C. Characterization of a glycerol/H+ symport in the halotolerant yeast Pichia sorbitophila. Yeast 1995;11:111–119.

44. Espeso EA, Tilburn J, Arst HN, Peñalva MA. pH regulation is a major determinant in expression of a fungal penicillin biosynthetic gene. EMBO J 1993; 12:3947–3956.

45. Ahuja M, Chiang Y-M, Chang S-L, Praseuth MB, Entwistle R, et al. Illuminating the diversity of aromatic Polyketide Synthases in *Aspergillus nidulans*. J Am Chem Soc 2012;134:8212–8221.

46. Khaldi N, Seifuddin FT, Turner G, Haft D, Nierman WC, et al. SMURF: Genomic mapping of fungal secondary metabolite clusters. Fungal Genet Biol 2010;47:736–741.

47. Oiartzabal-Arano E, Garzia A, Gorostidi A, Ugalde U, Espeso EA, et al. Beyond asexual development: Modifications in the gene expression profile caused by the absence of the *Aspergillus nidulans* transcription factor FlbB. Genetics 2015;199:1127–42.

48. Gerke J, Braus GH. Manipulation of fungal development as source of novel secondary metabolites for biotechnology. Appl Microbiol Biotechnol;98. Epub ahead of print 2014. DOI: 10.1007/s00253-014-5997-8.

49. Findon H, Calcagno-Pizarelli A-M, Martínez JL, Spielvogel A, Markina-lñarrairaegui A, et al. Analysis of a novel calcium auxotrophy in *Aspergillus nidulans*. Fungal Genet Biol 2010;47:647–655.

50. Shantappa S, Dhingra S, Hernández-Ortiz P, Espeso EA, Calvo AM. Role of the Zinc Finger Transcription Factor SltA in Morphogenesis and Sterigmatocystin Biosynthesis in the Fungus *Aspergillus nidulans*. PLoS One 2013;8:e68492.

51. Coker JA. Recent advances in understanding extremophiles [version 1; peer review: 2 approved]. F1000Research;8. Epub ahead of print 2019. DOI: 10.12688/f1000research.20765.1.

52. Gasch AP. Comparative genomics of the environmental stress response in ascomycete fungi. Yeast 2007;24:961–976.

53. Ianutsevich EA, Tereshina VM. Combinatorial impact of osmotic and heat shocks on the composition of membrane lipids and osmolytes in Aspergillus niger. Microbiology 2019;165:554–562.

54. Ni M, Yu J-H. A novel regulator couples sporogenesis and trehalose biogenesis in Aspergillus nidulans. PLoS One 2007;2:e970.

55. Kawasaki L, Sánchez O, Shiozaki K, Aguirre J. SakA MAP kinase is involved in stress signal transduction, sexual development and spore viability in Aspergillus nidulans. Mol Microbiol 2002;45:1153–1163.

56. Klein M, Swinnen S, Thevelein JM, Nevoigt E. Glycerol metabolism and transport in yeast and fungi: established knowledge and ambiguities. Environ Microbiol 2017;19:878–893.

57. Furukawa K, Hoshi Y, Maeda T, Nakajima T, Abe K. Aspergillus nidulans HOG pathway is activated only by two-component signalling pathway in response to osmotic stress. Mol Microbiol 2005;56:1246–1261.

58. Miskei M, Karányi Z, Pócsi I. Annotation of stress–response proteins in the aspergilli. Fungal Genet Biol 2009;46:S105–S120.

59. Harris SD, Turner G, Meyer V, Espeso EA, Specht T, et al. Morphology and development in *Aspergillus nidulans:* A complex puzzle. Fungal Genet Biol 2009;46:S82–S92.

60. Aranda A, del Olmo M l. Response to acetaldehyde stress in the yeast *Saccharomyces cerevisiae* involves a strain-dependent regulation of several ALD genes and is mediatedby the general stress response pathway. Yeast 2003;20:747–759.

61. Inglis DO, Binkley J, Skrzypek MS, Arnaud MB, Cerqueira GC, et al. Comprehensive annotation of secondary metabolite biosynthetic genes and gene clusters of *Aspergillus nidulans, A. fumigatus, A. niger* and *A. oryzae*. BMC Microbiol 2013;13:91.

62. Etxebeste O, Otamendi A, Garzia A, Espeso EA, Cortese MS. Rewiring of transcriptional networks as a major event leading to the diversity of asexual multicellularity in fungi. Crit Rev Microbiol 2019; 1–16.

