## Supplementary figures and images for "Defining the transcriptional responses of *Aspergillus nidulans* to cation/alkaline pH stress and the role of the transcription factor SltA"

### Figure S1

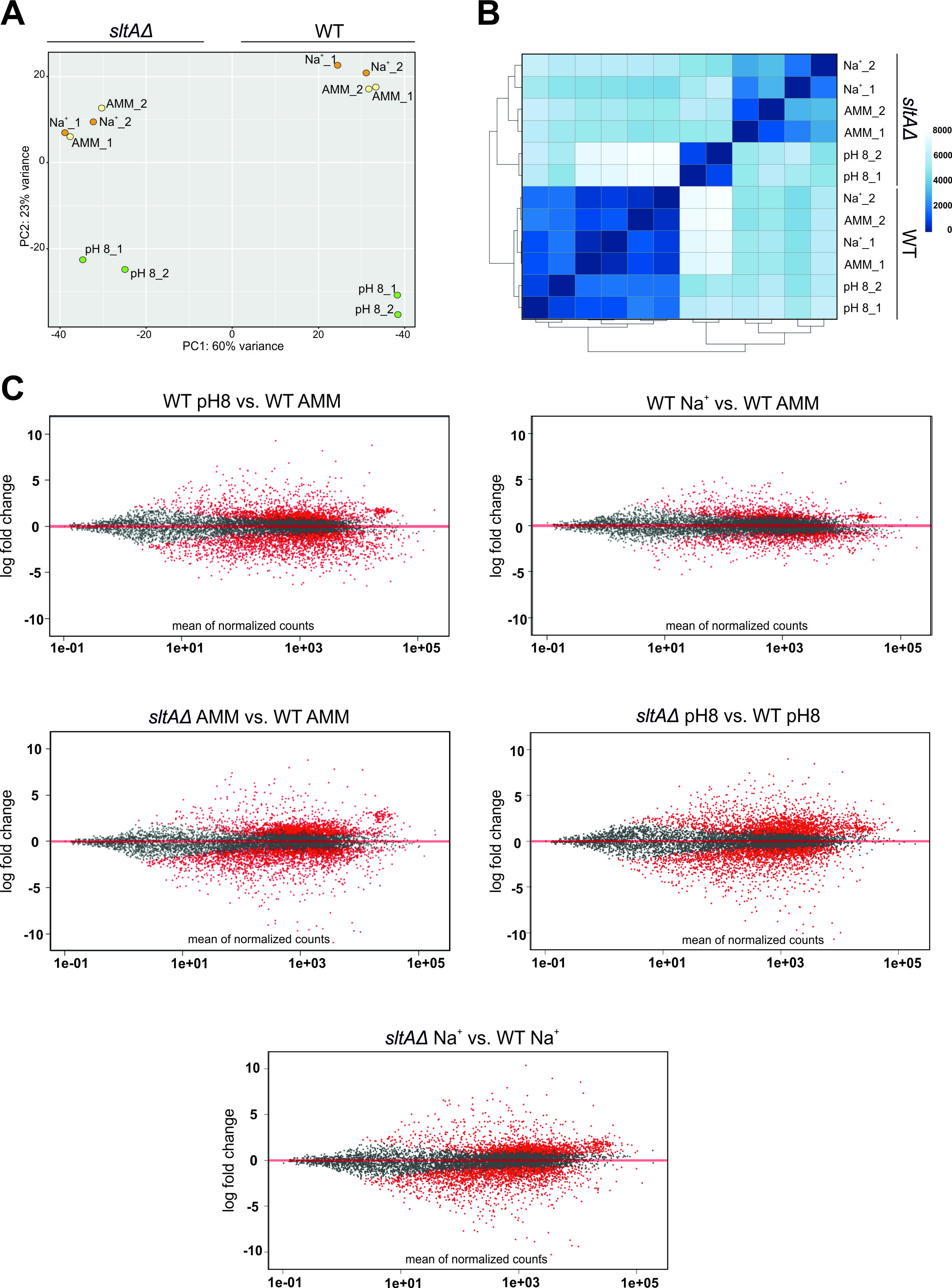

### Figure S2

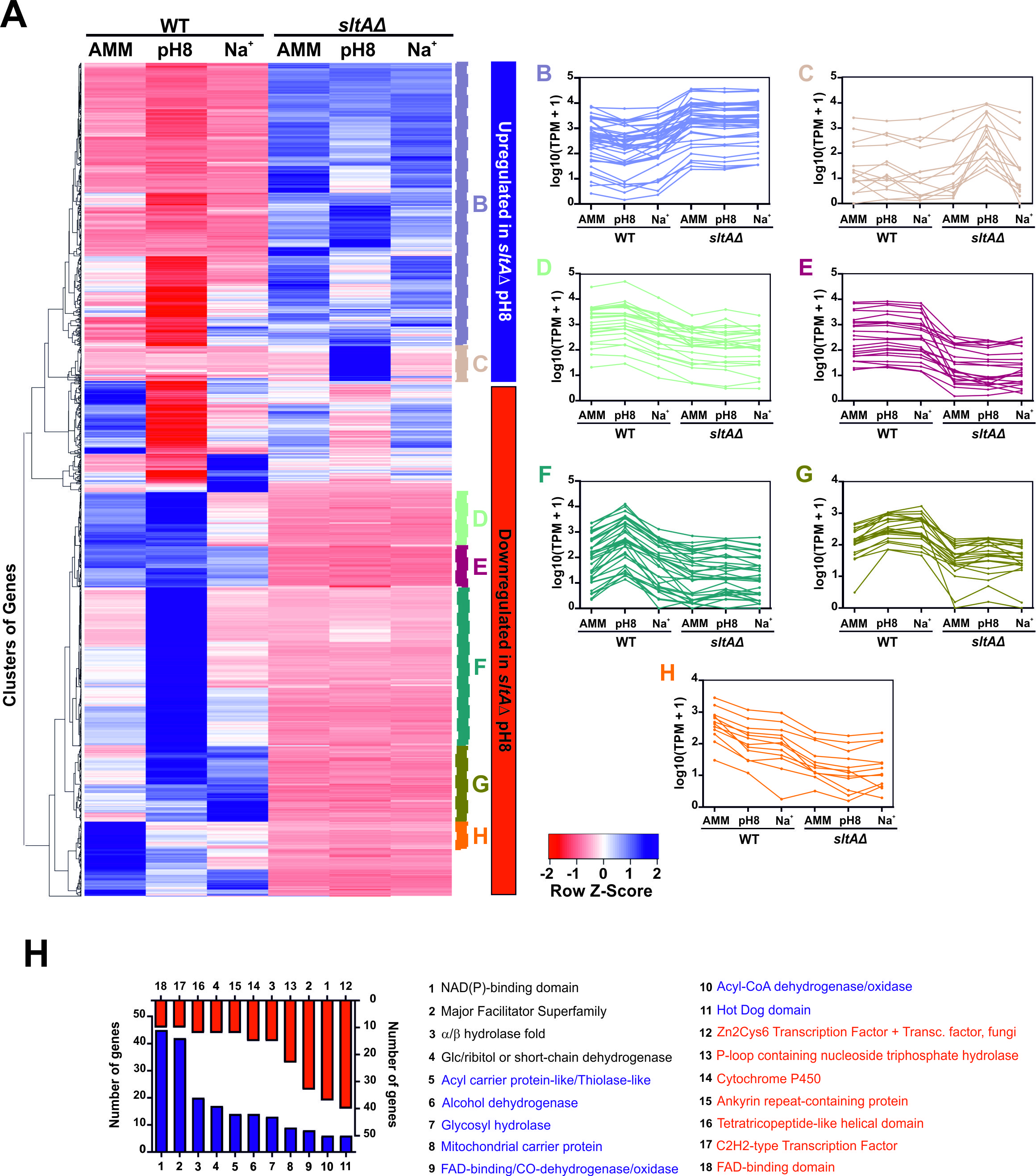

### Figure S3

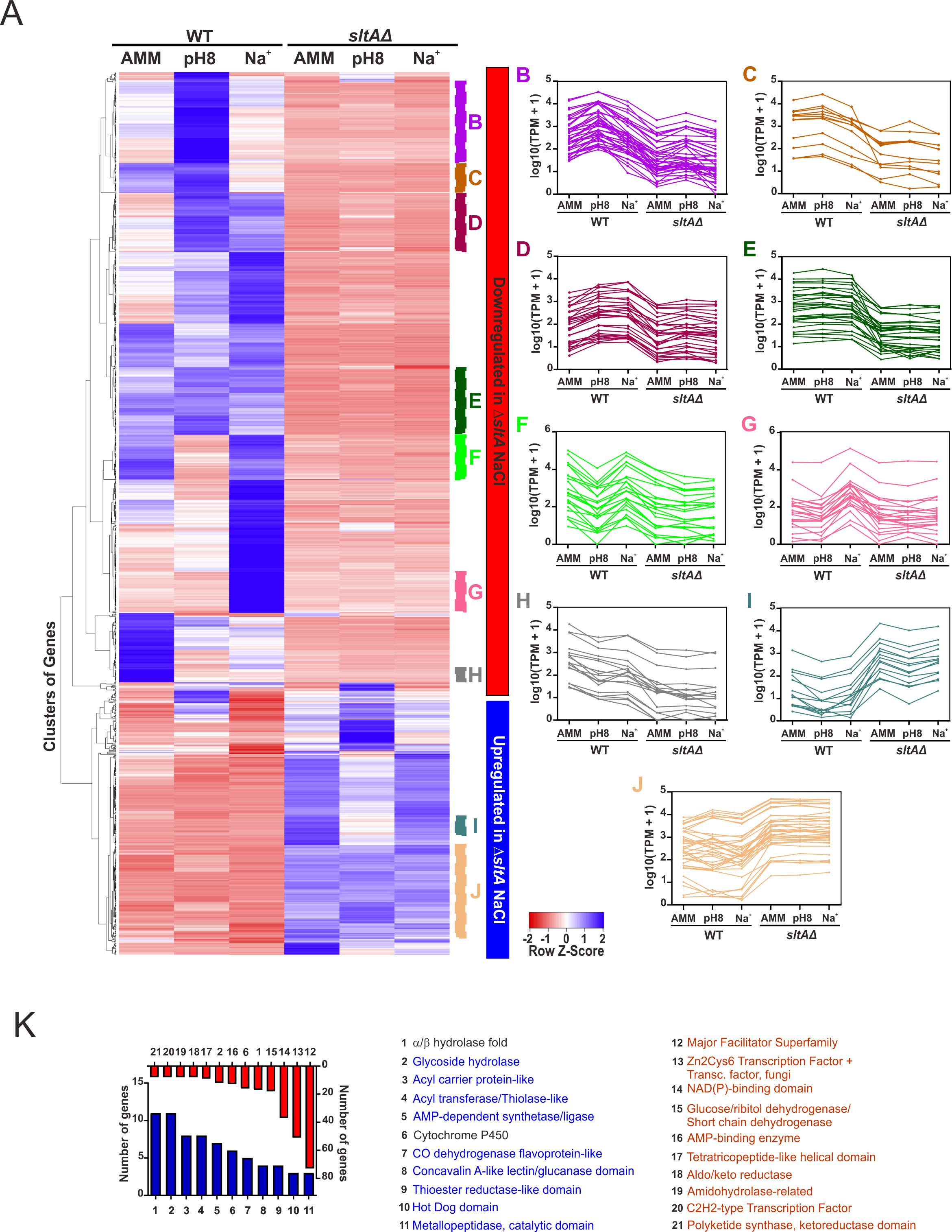
