## Supplementary material for "Defining the transcriptional responses of *Aspergillus nidulans* to cation/alkaline pH stress and the role of the transcription factor SltA": Table S1

**Supplementary Table S1**: Oligonucleotides used in this study.

| **Primer** | **Sequence (5'-3')** |
|---|---|
| qPCR-pacC1 | CTACATTGCCAACCGTCTTGAGC |
| qPCR-pacC2 | GGGATACATCACACTGTCCTCGG |
| qPCR-PalF1 | CTCTTGCCTAGTCAACCTCCGTG |
| qPCR-PalF2 | CGCTCCAACTCCTGTTTATCCTC |
| qPCR-BenA1 | AGATGCGCAACATCCAGAGC |
| qPCR-BenA2 | CGATCGTGGTACTCGGAGACCGA |
