## Supplementary material for "Defining the transcriptional responses of *Aspergillus nidulans* to cation/alkaline pH stress and the role of the transcription factor SltA": Table S2

**Supplementary Table S2**: Strains used in this study.

| Strain | Genotype | Reference |
|---|---|---|
| MAD4096 | *Wild type(prototroph)* | HHF27a, Findon et al., (2010) |
| MAD4097 | *sltAΔ::riboB*$^{Af}$ | HHF27b, Findon et al., (2010) |
| MAD1427 | *pyrG89, pabaB22,nkuAΔ::argB (argB2), riboB2* | TN02A25, Nayak et al.,(2006)) |
| MAD6669 | *pyrG89, pabaB22,nkuAΔ::argB (argB2), riboB2, sltA::ha*$_3$*::ribo*$^{Af}$ | This work |
| MAD3816 | pyroA4, nkuA*Δ::argB (argB2), sltAΔ::pyrG89, riboB2* | Mellado et al., (2016) |
| MAD3652 | *pyrG89, pabaB22, nkuAΔ::argB (argB2), sltA::ha*$_3$*::pyrG*$^{Af}$*, riboB2* | Mellado et al., (2016) |
